# METTL3-mediated m^6^A modification alleviates skeletal muscle atrophy in ALS by regulating ACSL1-dependent triglyceride metabolism

**DOI:** 10.64898/2026.10.21.759261

**Authors:** Qiupeng Yan, Lingyun Zhang, Xue Bai, Yingjun Guan, Haoyun Zhang, Jinmeng Liu, Baoxiang Zhuang, Mu Li, Xue Zhang, Xiaosu Chen, Xiaonan Ma, Chunjie Xu, Huilin Yang, Xin Huang, Fenghua Zhou, Yanchun Chen

## Abstract

Skeletal muscle-intrinsic defects are increasingly recognized as critical contributors to the pathogenesis of amyotrophic lateral sclerosis (ALS), yet the underlying molecular mechanisms remain poorly understood. *N*^6^-methyladenosine (m^6^A) is the most abundant post-transcriptional RNA modification in mammalian mRNAs that plays important regulatory roles in diverse physiological and pathological processes, however its role in ALS-associated skeletal muscle atrophy remains unknown. Here we found that global m^6^A methylation and the expression of its methyltransferase METTL3 are reduced in the skeletal muscle of hSOD1^G93A^ transgenic ALS mice. Multi-omics analyses reveal that m^6^A dysregulation is closely associated with impaired triglyceride metabolism. Functionally, METTL3 promotes triglyceride accumulation in ALS skeletal muscle in an m^6^A-dependent manner; overexpression of METTL3 in ALS myotubes improves energy supply and alleviates muscle atrophy by promoting triglyceride accumulation. *In vivo* overexpression of *Mettl3* in skeletal muscle attenuates muscle atrophy, improves motor performance, and prolongs survival of ALS mice. Mechanistically, we identify the fatty acid activation enzyme ACSL1 as a key downstream effector of METTL3-mediated regulation of triglyceride metabolism. METTL3-mediated m^6^A modification promotes the expression of *Acsl1* by enhancing its mRNA stability. Further investigation revealed that the regulatory effect of m^6^A on *Acsl1* expression depends on the m^6^A reader protein IGF2BP2. Collectively, our findings establish m^6^A modification as a critical regulator of skeletal muscle atrophy in ALS by maintaining triglyceride metabolic homeostasis, highlighting a promising therapeutic target for ALS.

## Introduction

Amyotrophic lateral sclerosis (ALS) is a fatal neurodegenerative disease characterized by progressive loss of motor neurons and widespread skeletal muscle atrophy. The disease typically exhibits an insidious onset and a rapidly progressive course, ultimately leading to muscle weakness, paralysis and respiratory failure. The median survival following symptom onset is approximately 2-4 years, reflecting its poor prognosis ^[1]^. Although numerous pathogenic genes, including *SOD1*, *C9ORF72*, and *FUS*, have been identified, the precise mechanisms underlying ALS pathogenesis remains unclear ^[2]^.

ALS has been viewed as a neurodegenerative disorder in which skeletal muscle atrophy is a secondary consequence of motor neuron loss ^[3]^. However, accumulating evidence supports the “dying-back” hypothesis, in which intrinsic skeletal muscle defects contribute to disease initiation and progression ^[4]^. Studies have shown that skeletal muscle is among the earliest structures affected in ALS, with pathological changes occurring even before significant motor neuron loss ^[5–9]^. Moreover, muscle-restricted expression of the pathogenic mutant hSOD1^G93A^ is sufficient to induce progressive muscle atrophy and motor dysfunction in mice, highlighting a potential muscle-autonomous role in ALS pathology ^[3, 5, 10]^. Nevertheless, the detailed intrinsic mechanisms driving skeletal muscle atrophy in ALS remain poorly understood.

Post-transcriptional RNA modifications have emerged as key regulators of gene expression and have been implicated in the development of various diseases ^[11]^. Among these, *N*^6^-methyladenosine (m^6^A) is the most abundant internal modification in mammalian mRNA ^[12]^. As a dynamic and reversible modification, m^6^A is catalyzed by the METTL3/METTL14/WTAP methyltransferase complex, in which METTL3 serves as the core catalytic component, and is removed by the demethylases FTO and ALKBH5 ^[13]^. m^6^A-mediated regulation of gene expression depends on reader proteins; for instance, YTHDF1 is known to control translation efficiency, whereas YTHDF2 and IGF2BP1/2/3 are involved in the regulation of mRNA stability ^[14, 15]^. Recent studies have reported that aberrant m^6^A modification contributes to motor neuron degeneration in ALS ^[16–21]^. However, whether m^6^A modification also participates in the regulatory mechanisms of intrinsic skeletal muscle atrophy in ALS remains unclear.

Skeletal muscle function is tightly coupled to metabolic homeostasis, particularly lipid metabolism, which serves as a major energy source through triglyceride storage and fatty acid oxidation. Clinically, patients with ALS frequently exhibit inadequate nutritional intake and a hypermetabolic state that accelerates energy expenditure, rendering skeletal muscle particularly vulnerable to energy imbalance ^[22]^. In this context, the maintenance of metabolic homeostasis likely requires dynamic regulation of gene expression, including post-transcriptional mechanisms such as m^6^A modification. Notably, emerging evidence indicates that m^6^A can modulate lipid metabolic pathways by regulating the expression of key enzymes. However, whether m^6^A-dependent control of lipid metabolism contributes to skeletal muscle atrophy in ALS remains unclear.

In this study, we identify a significant reduction of m^6^A modification levels and METTL3 expression in the skeletal muscle of ALS models. Through multi-omics analyses and experimental validation, we demonstrate that METTL3-mediated m^6^A modification alleviates ALS muscle atrophy by promoting triglycerides accumulation. Mechanistically, METTL3-mediated m^6^A promotes the expression of triglyceride biogenesis-related gene *Acsl1* by stabilizing its mRNA, and this regulatory effect is mediated by IGF2BP2. Our findings uncover a novel epitranscriptomic mechanism linking RNA methylation to lipid metabolism and muscle homeostasis, providing new insights into the muscle-autonomous pathogenesis of ALS.

## Materials and Methods

### Animals

hSOD1^G93A^-tg mice (ALS mice) and wild-type mice (WT mice) were purchased from Jackson Laboratory (Bar Harbor, Maine, USA). Animals were born and raised in a specific-pathogen-free environment and on a standard chow diet under a temperature of 23±□1□°C, a humidity of 45% and a 12 h light-dark cycle. All mice had access to food and water *ad libitum*. All mice were group caged by sex at adolescence, with 3-5 individuals for each cage. Genotype of each mouse was determined by PCR based on the tail tissue derived genomic DNA. Unless otherwise specified, both male and female mice were randomly assigned to experimental groups. All experiments were approved by the Medical Ethics Committee of Shandong Second Medical University (protocol code 2024SDL446) and were conducted in accordance with the guidelines of the Helsinki Declaration

### Behavioral

#### General conditions of behavioral tests

All behavioral tests were carried out on mice at the age of 9-15 weeks. All tests were conducted only during the light cycle (07:00 ∼ 19:00). For animal habituation, all mice were first handled ∼2 min for 3 successive days and transferred to the testing room at least 12 h prior to behavioral tests. Mice subjected to different behavioral tests had at least an interval of 2 h between two tests.

#### Rotarod test

Animals (5 per group) were gently placed on a rotating rod (Med Associates Inc., USA) at an initial speed of 2 r/min for 30 s of habituation. Then, the rotating speed gradually accelerated from 2 r/min to 20 r/min in 300 s. The time spent by animals on the rotarod was recorded by electric pedal when animals fell down.

#### Grip strength

Only male mice were subjected to grip strength test. Animals were placed on the grip force meter (Shenzhen Ruiwode Life Technology, China) and waiting for stable. During measurements, pull the mouse body backward with constant speed until the mouse four paws release, and the maximum grip force was recorded. Each mouse is measured three times, and the maximum value is used for analysis.

#### Walking gait analysis

The limbs of animals were marked with non-toxic, water-soluble paint in distinct colors (red for forelimbs and blue for hindlimbs). Mice were then placed at one end of a 40 cm × 5 cm homemade walkway with side barriers and allowed to walk to the other end without disturbance. Paw prints were recorded for analysis. Step length (distance between consecutive paw center of the same limb) was measured for both forelimbs and hindlimbs.

#### Cell culture

C2C12 myoblasts were purchased from ATCC and maintained in Dulbecco’s Modified Eagle Medium (DMEM) supplemented with 10% fetal bovine serum (FBS; Gibco, Rockville, MD, USA), 100 U/mL penicillin, and 100 µg/mL streptomycin at 37 °C in a humidified atmosphere containing 5% CO . Mycoplasma contamination was routinely tested on a weekly basis. For transfection, cells were transfected using Lipofectamine 2000 (Invitrogen, Carlsbad, CA, USA) according to the manufacturer’s instructions. For differentiation, the culture medium was replaced with DMEM containing 2% horse serum. Cells were maintained under these conditions for 96 h, with the medium refreshed every 48 h. Sequence of small interfering RNAs (siRNAs) used in this article were provided in Supplementary Data 5

#### Energy deprivation experiments

C2C12 myoblasts were first transfected and differentiated into myotubes as described above. After 96 h of differentiation, the differentiation medium was removed, and cells were washed twice with PBS and incubated in D-Hank’s balanced salt solution for 4 h. Cell survival was then assessed using the CCK-8 assay.

#### Triglyceride assay

Triglyceride levels in skeletal muscle tissue and myotubes were measured using the Amplex Red TG Assay Kit (Beyotime, China) according to the manufacturer’s instructions. For skeletal muscle, tissue samples were weighed and homogenized in 1 mL isopropanol per gram of tissue, followed by centrifugation at 12,000 rpm for 10 min at 4 °C. The supernatant was collected for analysis and triglyceride concentration was normalized by tissue weight. For myotubes, cells were harvested with trypsin, counted, and lysed in 200 μL isopropanol per 10 cells. The lysates were sonicated and centrifuged at 12,000 rpm for 10 min at 4 °C, and the resulting supernatants were used for detection.

#### Fatty acid oxidation assay

Fatty acid oxidation activity was assessed using the FAO Blue reagent (Funakoshi, Japan) according to the manufacturer’s protocol. Briefly, the differentiation medium was removed, and cells were washed twice with PBS and incubated in serum- and phenol red-free DMEM. FAO Blue was then added at a final concentration of 20 μM, and cells were incubated for 2 h. Blue fluorescence intensity was measured at 480 nm using SpectraMax iD3 (Molecular Devices) fluorescence microplate reader.

#### ATP content assay

ATP levels in myotubes were measured using the ATP Assay Kit (Byotime, China) according to the manufacturer’s instructions. Briefly, cells were harvested with trypsin, counted, and lysed in 200 μL lysis buffer per 10 cells. The lysates were sonicated and centrifuged at 12,000 rpm for 10 min at 4 °C, and the resulting supernatants were used for ATP detection.

#### Antibodies and chemicals

Anti-m^6^A (Zenbio, Cat. no. 517924, 1:1000), Anti-METTL3 (Proteintech, Cat. no. 15073-1-AP, 1:1000), Anti-IGF2BP2 (Proteintech, Cat. no. 11601-1-AP), Anti-GAPDH (Proteintech, 60004-1-Ig, 1:20000), FITC-labeled α-Bungarotoxin (α-BTX) (MedChemExpress, Cat. no. HY-P1264F, 1:200), STM2457 (MedChemExpress, Cat. no. HY-134836, 25 μM), T863 (MedChemExpress, Cat. no. HY-32219, 10 μM).

#### H&E staining

Mouse tissues were fixed in 10% neutral-buffered formalin, dehydrated, cleared, and embedded in paraffin. Paraffin blocks were cooled on ice or at 4 °C for 20 min and sectioned at 4 µm thickness. Sections were mounted on glass slides and baked at 60 °C for 2 h. Slides were deparaffinized twice in xylene (15 min each), followed by rehydration through a graded ethanol series (100%, 95%, 80%, and 70%) and rinsed in water to achieve full hydration. Sections were stained with hematoxylin for 10 min, rinsed in running water, and differentiated in 1% acid alcohol for 20 s. After rinsing, slides were blued in 0.1% ammonia water for 2 s and washed for 10 min. Counterstaining was performed with 0.5% eosin for 1 min, followed by rinse in water and dehydration through graded ethanol (70%-100%). Finally, sections were cleared twice in xylene (15 min each) and mounted with neutral resin.

#### Oil Red O staining

Frozen gastrocnemius samples were embedded in OCT compound and cryosectioned at a thickness of 12 µm. Sections were washed with PBS for 20 s and stained with modified Oil Red O solution (Beyotime, China) for 30 min at room temperature in the dark. After staining, slides were briefly rinsed with differentiation solution for 30 s, washed thoroughly with PBS for 20 s, and mounted with an antifade mounting medium (Beyotime, China). Images were acquired under a light microscope.

#### Immunofluorescence

Differentiated C2C12 myotubes were washed twice with PBS and fixed with 4% paraformaldehyde at 4 °C for 15 min. After three PBS washes, cells were permeabilized with 0.3% Triton X-100 for 10 min at room temperature and washed again. Blocking was performed with 10% goat serum at 37 °C for 30 min. Cells were then incubated overnight at 4 °C with rabbit anti-MYHC antibody (1:200), followed by FITC-conjugated goat anti-rabbit IgG (1:200) for 50 min at 37 °C. After three PBS washes, cells were mounted for fluorescence microscopy.

#### BODIPY staining

Differentiated C2C12 myotubes were washed twice with PBS and fixed with 4% paraformaldehyde at 4 °C for 15 min. After three washes with PBS, cells were blocked with 10% goat serum at 37 °C for 30 min, followed by incubation with BODIPY (1:1000) and DAPI (1:800) for 10 min at 37 °C. Following three additional PBS washes, cells were either maintained in PBS for fluorescence microplate analysis or mounted for fluorescence microscopy.

#### WGA staining

WGA staining was performed using the iFluor 488-WGA staining kit (Shanghai Huilan Biological Technology, China) according to the manufacturer’s instructions with minor modifications. Briefly, frozen sections of skeletal muscle were subjected to antigen retrieval in sodium citrate buffer (pH 6.0), permeabilized with 0.3% Triton X-100, and blocked with 10% goat serum at room temperature. Sections were then incubated with iFluor 488-WGA at 37 °C for 2 h. After washing with PBS, sections were mounted for fluorescence microscope.

#### α-BTX staining

Frozen sections of mouse skeletal muscle were subjected to antigen retrieval in sodium citrate buffer (pH 6.0). Sections were then permeabilized with 0.3% Triton X-100 and blocked with 10% goat serum at room temperature, followed by incubation with FITC-conjugated α-BTX at 37 °C for 1 h. After washing with PBS, sections were mounted and imaged under a fluorescence microscope.

#### Protein extraction and immunoblotting

For protein extraction from tissue samples, animal tissues were first frozen in liquid nitrogen, ground into powder and lysed in RIPA (Beyotime, China, P0013B). For cultured cell samples, cells were washed with cold PBS and collected with cell scraper in RIPA. The protein lysis was sonicated, centrifuged and the supernatant was denatured in SDS-loading buffer (Genestar). Protein concentration was assayed by BCA. Equal amounts of protein samples were separated on SDS-PAGE gel and transferred onto a 0.22-μm PVDF membrane in wet-transfer system (BioRad). Then the membranes were firstly blocked with 5% skim milk at room temperature for 2 h, incubated with primary antibodies at 4 °C overnight and incubated with secondary antibodies at room temperature for 1 h. The membranes were washed with 1×TBST (5 times × 1 min) after each incubation process. Protein signal was detected with the ChemiDoc^TM^ Imaging Systems (BIO-RAD).

#### RNA isolation

Each tissue or cell sample was lysed in 1 ml M5 SuperPure Total RNA Extraction Reagent (Mei5, China), followed by chloroform (200 μL) extraction and isopropanol (500 μL) precipitation to isolate total RNA. The concentration of RNA was measured by NANODROP 2000c spectrophotometer (Thermo Fisher Scientific). Only samples with OD 260/280 nm ratio more than 1.8 were used in subsequent experiments.

#### Untargeted lipidomics

Skeletal muscle samples were homogenized in liquid nitrogen and sonicated in 400 µL cold 75% methanol for 15 min. Plasma and urine were processed directly. MTBE (1 mL) was added and shaken for 1 h at room temperature. Phase separation was induced with 250 µL water, followed by incubation for 10 min and centrifugation (14,000 g, 15 min, 4□). The upper organic phase was collected and dried under nitrogen. QC samples were generated by pooling aliquots from all samples. Dried extracts were resuspended in 50% acetonitrile, filtered (0.22 µm), transferred to HPLC vials, and stored at -80□. Samples were analyzed using a Q Exactive Plus Orbitrap coupled to a Nexera X2 UHPLC with a Hypersil GOLD C18 column (100 × 2.1 mm, 1.9 µm). Mobile phases consisted of ammonium acetate/acetonitrile (A) and acetonitrile/isopropanol (B). A 30–100% B gradient was used at 0.3 mL/min. Data were acquired in positive and negative ESI modes using data-dependent Top10 fragmentation. Raw data were processed in MS-DIAL (LipidBlast library). Tolerances were 10 ppm (precursor) and 20 ppm (fragment). Only lipid features with <30% RSD in QC samples were retained. Untargeted lipidomics analysis was conducted by Shanghai Bioprofile Technology Company Ltd.

#### RT-qPCR

RNA samples were firstly treated with DNase I to remove genomic DNA, then reverse transcribed into cDNA by reverse transcriptase (ThermoFisher Scientific). Oligo dT (Promega) was used as primers during reverse transcription reaction. SYBR Green Mix (Mei5) was used in qPCR experiments, and relative gene expression was normalized to *Rps18* with the 2^−△△Ct^ method. Primers used in this article were provided in Supplementary Data 5.

#### Methylated RNA dot blot assay

For the dot blot assay, 1 μg of total RNA was spotted onto a positively charged nylon membrane. Membranes were blocked with 5% skim milk for 2 h at room temperature, followed by incubation with an anti-m^6^A primary antibody overnight at 4 °C. After washing with 1× TBST (5 times × 1 min), membranes were incubated with an appropriate HRP-conjugated secondary antibody for 2 h at room temperature and washed again with 1× TBST (5 times × 1 min). Signals were detected using the ChemiDoc^TM^ Imaging Systems (BIO-RAD).

#### RNA immunoprecipitation (RIP)

RNA immunoprecipitation was performed using the PureBinding RNA Immunoprecipitation Kit (Geneseed) according to the manufacturer’s instructions. Briefly, skeletal muscle tissue was weighed and lysed with RIP lysis buffer (100 mg tissue/mL). The 400 μL supernatant was then incubated overnight at 4 °C with A/G beads coupled to either IGFBP2 antibody or control IgG (5 μg antibody per reaction). Meanwhile, 100 μL of the supernatant was set aside as input controls for RNA extraction. The immunoprecipitated RNA and input samples were subsequently analyzed by RT qPCR.

#### Methylated RNA immunoprecipitation (MeRIP)

Methylated RNA immunoprecipitation was performed using the EpiQuik CUT&RUN m^6^A RNA Enrichment Kit (EpigenTek, USA) with minor modifications. Briefly, 20 μg of total RNA was incubated with anti-m^6^A antibody or non-immune IgG-coupled A/G beads for 90 min; IgG served as the negative control. CUT&RUN was then performed using the Cleavage Enzyme Mix for 4 min followed by immunoprecipitation. In parallel, 500 ng of non-immunoprecipitated input RNA was incubated with the Cleavage Enzyme Mix for 4 min. Immunoprecipitated RNA and input RNA were purified using the ZYMO RNA Clean & Concentrator Kit (Zymo Research, USA) according to the manufacturer’s instructions. RNA quality and fragment size distribution were analyzed using the Agilent 2100 Bioanalyzer, and samples with RNA fragments of ∼200 bp were used for subsequent experiments.

#### RNA-seq analysis

RNA-seq data were obtained from the GEO database under project IDs GSE245520 ^[23]^ and GSE215424 ^[24]^. Raw FASTQ files were downloaded and processed using Fastp (version 0.23.4) for quality control and adapter trimming ^[25]^. Clean reads were aligned to the reference genomes (GRCm38, Ensembl release 84 for mouse; GRCh38, Ensembl release 95 for human) using HISAT2 (version 2.2.0) with default parameters ^[26]^. Aligned reads were quantified using StringTie (version 2.2.1) with parameters -e-B. Read count matrices were generated with the prepDE.py script provided by StringTie ^[27]^. Differential gene expression analysis was performed using the DESeq2 (version 1.34.0) package in R ^[28]^. Functional enrichment analysis was conducted using the Metascape online platform (https://metascape.org/) ^[29]^, and gene set enrichment analysis (GSEA) was performed with the ClusterProfiler (version 4.2.2) package in R ^[30]^.

#### m^6^A-MeRIP-seq analysis

m^6^A-MeRIP-seq data were obtained from the GEO database under project ID GSE179368 ^[31]^. Raw FASTQ files were downloaded and subjected to adapter and quality trimming using Fastp (version 0.23.4) ^[25]^. Clean reads were aligned to the mouse reference genome (GRCm38, Ensembl release 84) using HISAT2 (version 2.2.0) with default parameters ^[26]^. m^6^A modification peaks were identified using the exomePeak2 (version 1.6.1) package in R ^[32]^. The screening criteria for m^6^A peaks were defined as follows: Log_2_(IP/Input) fold change > 2, FDR < 0.05, RPM(IP) > 0.2, and only peaks located within protein-coding genes were retained. Normalized coverage tracks were generated with DeepTools bamCoverage (version 3.5.5) with parameters --binSize 10 --normalizeUsing RPKM for visualization of m^6^A peak distribution in Integrative Genomics Viewer (version 2.11.2).

#### Data statistics and visualization

Unless stated otherwise, unpaired two-tailed Student’s *t*-test was used for comparison between two groups, and one-way ANOVA was used for three or more groups comparison. All data visualization was generated from Graphpad prism (version 10.0) or R package ggplot2 (Version 3.3.6). The schematic diagrams was drawn by the GDP online website ^[33]^. The error bars represent mean ± SEM.

## Results

### m^6^A modification and METTL3 expression are decreased in ALS skeletal muscle

To characterize the dynamic profile of m^6^A modification in skeletal muscle during disease progression, we quantified global m^6^A levels in the skeletal muscle of hSOD1^G93A^ transgenic (hSOD1^G93A^-tg) ALS mice at different disease stages (Fig. 1A) ^[34, 35]^. Decreased m^6^A modification levels were detected in the skeletal muscle of ALS mice compared with wild-type controls (WT). Notably, this decrease of m^6^A levels was already detectable at the pre-symptomatic stage (Fig. 1A, B), indicating that dysregulation of m^6^A modification represents an early molecular event in ALS pathogenesis.

**Figure 1.**
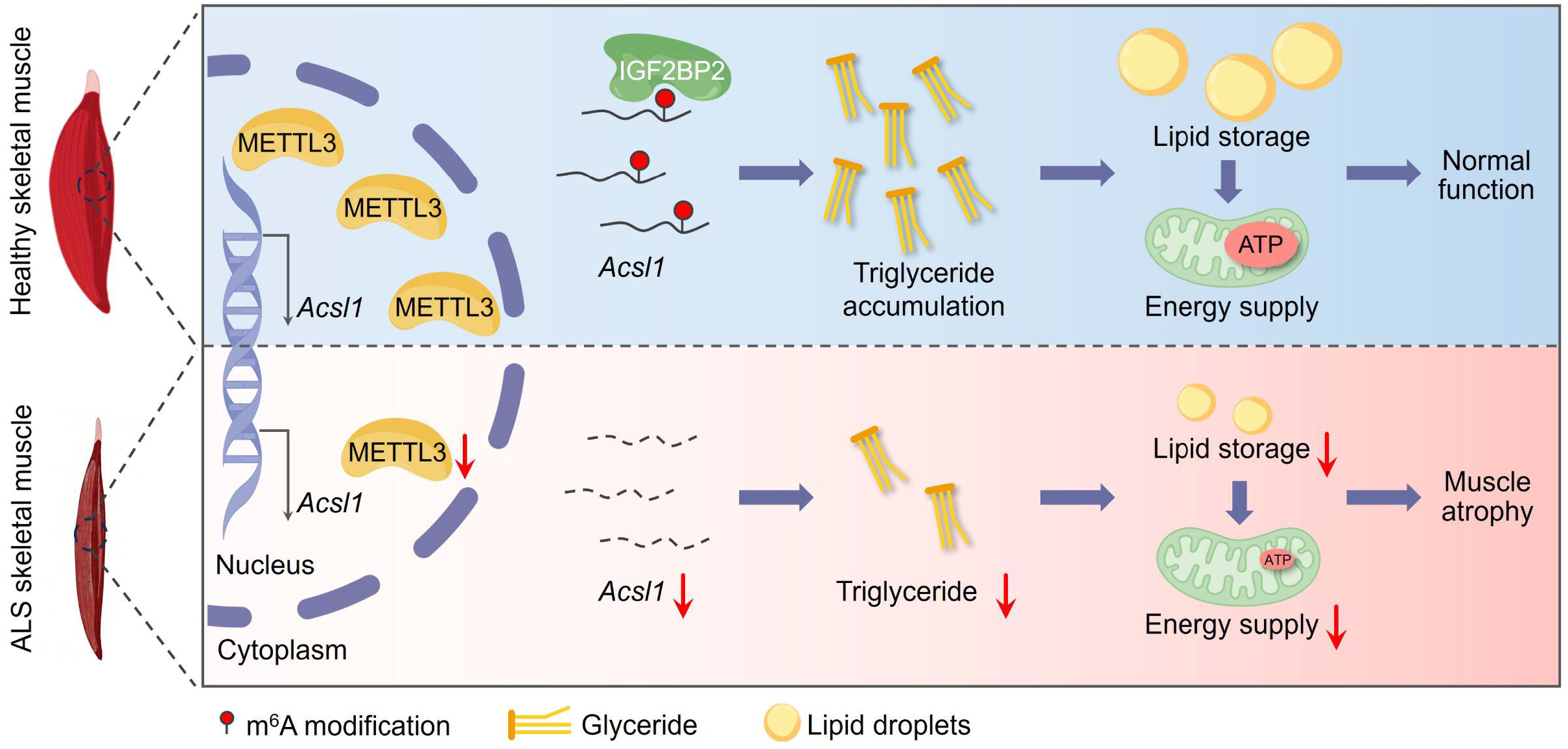
Reduced m^6^A modification levels and METTL3 expression in the skeletal muscle of hSOD1^G93A^ ALS mice. **(A)** Schematic diagram of the hSOD1^G93A^-tg ALS mouse model. This model shows no evident neuronal loss or symptoms at early stages. Clinical symptoms typically appear at ∼13 weeks of age and progress rapidly over the following 2 weeks. By ∼15 weeks, mice exhibit severe limb motor impairment, with an average lifespan of ∼18 weeks. **(B)** Dot blot analysis of global m^6^A modification level in the gastrocnemius of WT and hSOD1^G93A^-tg ALS mice at indicated time point (n = 3 per group). **(C)** RT-qPCR analysis of mRNA expression of m^6^A writer genes (*Mettl3*, *Mettl14* and *Watp*) and eraser genes (*Fto* and *Alkbh5*) in gastrocnemius of WT and hSOD1^G93A^-tg ALS mice at age of 15 weeks (n = 3 per group). **(D)** RT-qPCR analysis of *Mettl3* mRNA expression in WT and hSOD1^G93A^-tg ALS mice at indicated time point (n = 3 per group). **(E)** Western blot analysis of METTL3 protein expression in gastrocnemius of WT and hSOD1^G93A^-tg ALS mice at indicated time points (n = 3 per group) Data are presented as mean ± SEM. \**P* < 0.05, \*\**P* < 0.01, \*\*\**P* < 0.001; n.s., not significant.

To identify the upstream regulators responsible for m^6^A alteration, we assessed the expression of key m^6^A writers and erasers ^[13]^. Results showed that *Mettl3* expression was significantly reduced in the skeletal muscle of ALS mice, while other writer genes (*Mettl14* and *Wtap*) or eraser genes (*Fto* and *Alkbh5*) remained unchanged (Fig. 1C). In line with reduced global m^6^A levels, *Mettl3* mRNA and protein expression were already decreased at the pre-symptomatic stage (Fig. 1D, E). Collectively, these findings demonstrate that m^6^A modification and METTL3 expression are dysregulated in ALS skeletal muscle, suggesting a close association with muscle atrophy in ALS.

### Impaired METTL3-mediated m^6^A modification contributes to triglyceride downregulation in ALS skeletal muscle

To elucidate the biological processes potentially regulated by m^6^A in ALS skeletal muscle, we first analyzed the m^6^A epitranscriptome of mouse skeletal muscle using publicly available datasets (Supplementary Data 1) ^[31]^. Functional enrichment analysis revealed that m^6^A-modified genes were associated with lipid metabolism-related processes, supporting a close association between m^6^A modification and lipid metabolic regulation (Fig. S1A, B, Supplementary Data 2).

Given the critical role of m^6^A in post-transcriptional gene regulation ^[13]^, we hypothesized that lipid metabolism-related genes are dysregulated in the skeletal muscle of ALS models. To test this hypothesis, we further analyzed skeletal muscle RNA-seq data of ALS mice obtained from public datasets (Fig. 2A) ^[23]^. Principal component analysis (PCA) showed a clear separation between ALS and WT samples (Fig. 2B). Functional enrichment analysis indicated that the downregulated genes were significantly enriched in lipid metabolism-related pathways (Fig. 2C-F, Supplementary Data 3). Consistently, gene set enrichment analysis (GSEA) demonstrated suppression of multiple lipid biosynthesis and catabolism pathways in ALS skeletal muscle (Fig. 2G). To assess whether these changes are conserved in humans, we further examined skeletal muscle RNA-seq data of ALS patients from public datasets (Fig. 2H) ^[24]^. PCA again revealed distinct transcriptomic profiles between ALS and healthy controls (Fig. 2I), and GSEA confirmed significant downregulation of lipid biosynthesis- and catabolism-related pathways (Fig. 2J). These results indicate that lipid metabolism-related genes are suppressed in the skeletal muscle of both ALS mice and patients.

**Figure 2.**
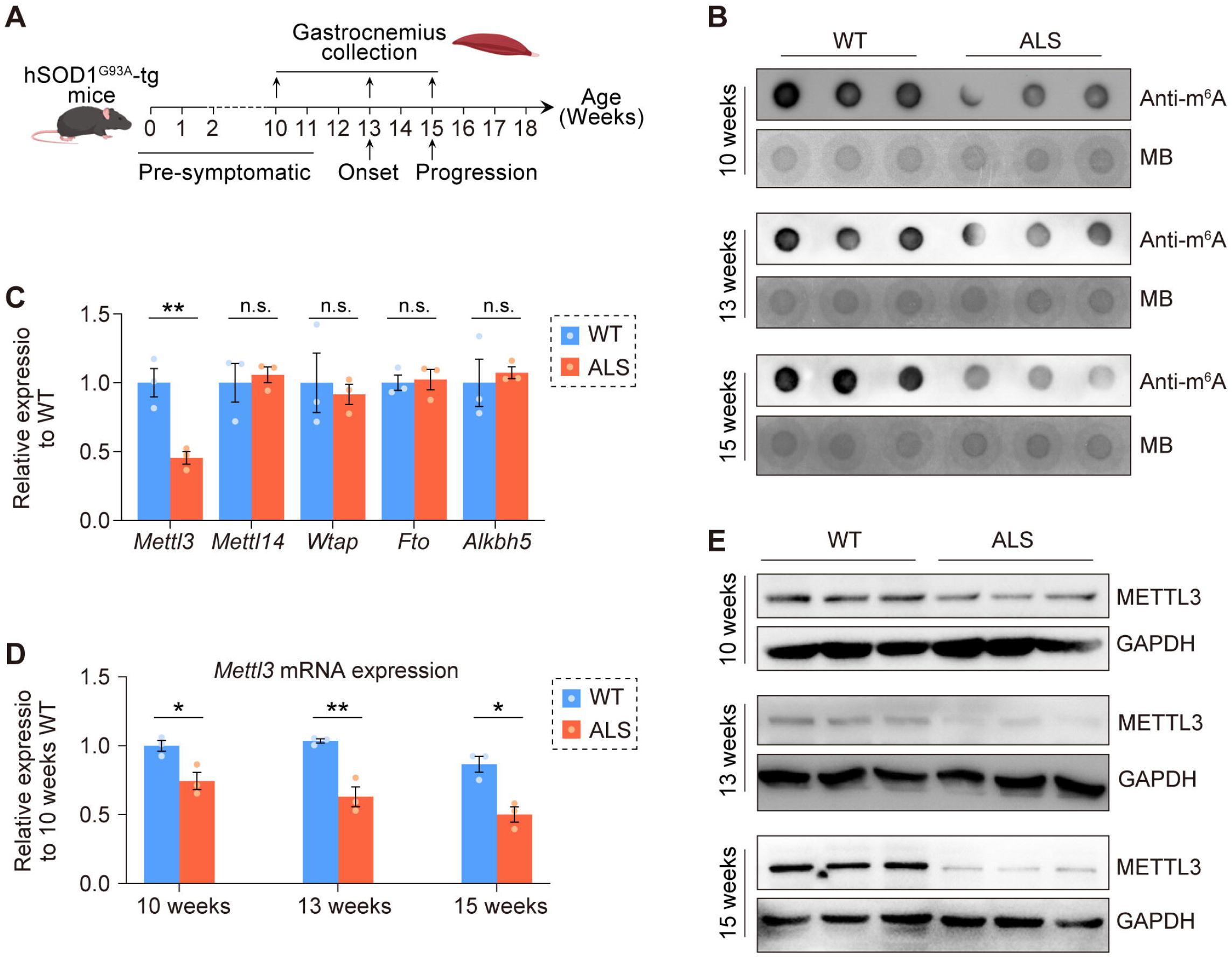
Lipid metabolism-related genes dysregulated in ALS skeletal muscle. **(A)** Schematic overview of publicly available RNA-seq datasets of tibialis anterior from ALS mice (*Fus*Δ*14* mutation) (GEO accession: GSE245520). **(B)** PCA of RNA-seq data from tibialis anterior of WT and ALS mice. **(C)** Volcano plot showing differentially expressed genes (DEGs) between tibialis anterior of WT and ALS mice. DEGs were defined by fold change > 1.25 and *P* < 0.05. **(D)** Functional enrichment analysis of downregulated genes in tibialis anterior of ALS mice. **(E)** Network visualization of enriched functional terms. Each circle represents a biological process, with node size proportional to the number of associated genes, edge thickness indicating similarity score, and color representing adjusted *P*-value. **(F)** Heatmap of lipid metabolism-related DEGs in WT and ALS tibialis anterior. Genes were annotated according to GO term “lipid metabolic process” (GO:0006629). **(G)** Gene set enrichment analysis (GSEA) of all expressed genes in WT and ALS tibialis anterior. **(H)** Schematic overview of publicly available RNA-seq datasets of skeletal muscle from ALS patients (GEO accession: GSE215424). **(I)** PCA of RNA-seq samples from skeletal muscle of ALS patients and healthy controls. **(J)** GSEA of all expressed genes in skeletal muscle of ALS patients and healthy controls.

To further characterize detailed lipid metabolic disturbances, we performed untargeted lipidomics analysis in gastrocnemius of the hSOD1^G93A^ transgenic (hSOD1^G93A^-tg) ALS mice (Fig. S2A). PCA revealed a clear separation between ALS and WT samples, indicating a distinct alteration in the lipid metabolic landscape (Fig. 3A). Differential metabolite analysis revealed 60 upregulated and 109 downregulated lipid species (Fig. S2B, Supplementary Data 4). Classification of these differential metabolites by lipid class demonstrated that fatty acyls (FA) and glycerolipids (GL) exhibited a high proportion of downregulated species, with 100% and 93% of differential metabolites being downregulated, respectively (Fig. 3B, C). Within the FA class, the downregulated metabolites were primarily confined to the N-acyl amino acids (NAR) and carnitine esters (CAE), both of which are present at relatively low abundance, suggesting a limited contribution to global lipid remodeling (Fig. S2C, Fig. 3D). However, glycerolipids accounted for 5 of the top 10 contributors identified by PCA loading analysis, highlighting their central role in driving lipidomic divergence between WT and ALS skeletal muscle. (Fig. S2D). Within the glycerolipid class, triglycerides constituted the most abundant lipid species across all detected lipids, and notably, all differential triglyceride species were downregulated (Fig. 3D, E). Given that triglycerides serve as the key reservoir for energy storage in skeletal muscle, their consistent downregulation likely reflects impaired lipid storage capacity and disrupted energy homeostasis ^[36]^. These findings identify triglycerides as a key lipid class underlying metabolic remodeling associated with skeletal muscle atrophy in ALS.

**Figure 3.**
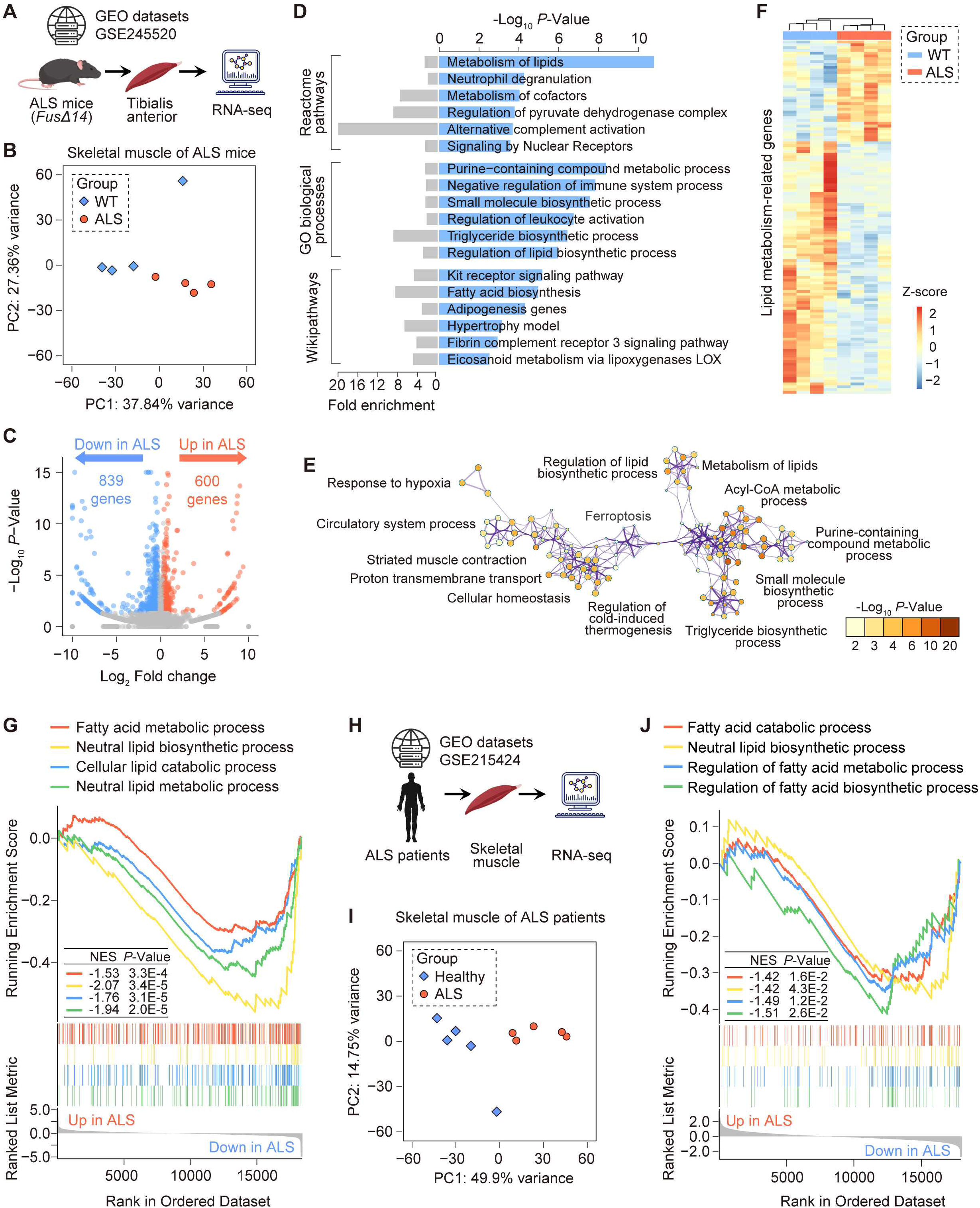
Triglyceride levels decreased in the skeletal muscle of hSOD1^G93A^-tg ALS mice. **(A)** PCA of lipidomics profiles from gastrocnemius of WT and hSOD1^G93A^-tg ALS mice **(B)** Scatter plot of lipid metabolites categorized by lipid class. Each point represents a lipid metabolite; point size indicates Variable importance in projection (VIP) value and color denotes lipid category. FA, fatty acyl; GL, glyceride; GP, glycerophospholipid; ST, sterol; Unknown, unidentified lipid species. **(C)** Stacked bar chart showing the proportion of upregulated and downregulated lipid metabolites among total differential metabolites. **(D)** Proportion of molecular counts across different lipid subclasses in gastrocnemius of WT and hSOD1^G93A^-tg ALS mice. Triglyceride was labeled by red color. **(E)** Heatmap displaying changes in triglyceride species between the gastrocnemius of WT and hSOD1^G93A^-tg ALS mice. Triglycerides was labeled by red color. **(F)** Amplex red assay of triglyceride (n = 4 per group) levels in gastrocnemius of hSOD1^G93A^-tg ALS mice at indicated time point. **(G)** Oil Red O staining of gastrocnemius from hSOD1^G93A^-tg ALS mice at indicated time point. Scale bar represents 100 μm. **(H)** Quantification of Oil Red O staining intensity from panel **(F)** (n (10 weeks) = 8 (WT) and 8 (ALS), n (14 weeks) = 7 (WT) and 6 (ALS), n (15 weeks) = 8(WT) and 8(ALS)). **(I)** Amplex red assay of triglyceride levels in hSOD1^G93A^-tg ALS myotubes after *Mettl3* overexpression (n = 4 per group). **(J)** Representative BODIPY staining of hSOD1^G93A^-tg ALS myotubes after *Mettl3* overexpression. Scale bar represents 100 μm. **(K)** Quantification of BODIPY fluorescence intensity. Data are presented as mean ± SEM. \**P* < 0.05, \*\*\**P* < 0.001; n.s., not significant.

We then validated triglyceride alterations in the gastrocnemius of hSOD1^G93A^-tg ALS mice at different disease stages, results showed that triglyceride levels were already reduced at 10 weeks of age, time point before the onset of clinical symptoms, and remained decreased at 15 weeks old, when mice exhibited severe motor dysfunction (Fig. 3F). Oil Red O staining further confirmed a reduction in neutral lipid content (Fig. 3G, H). Together, our lipidomics analysis and experiments demonstrate that triglycerides are diminished in ALS skeletal muscle.

To further investigate the functional role of m^6^A modification in triglyceride metabolism, we overexpressed *Mettl3* in an *in vitro* ALS myotube model, in which hSOD1^G93A^ expressing C2C12 myoblasts were differentiated into myotubes (Fig. S3A-D). Results showed that ALS myotubes exhibited significantly reduced triglyceride level, whereas *Mettl3* overexpression restored triglyceride abundance (Fig. 3I). In contrast, overexpression of a catalytically inactive *Mettl3* mutant (*Mettl3^Mut^*) failed to rescue triglyceride levels (Fig. 3I), indicating that the regulatory effect of METTL3 on triglyceride metabolism is dependent on its methyltransferase activity. BODIPY staining further confirmed that *Mettl3*, but not *Mettl3^Mut^*, promotes neutral lipid accumulation in ALS myotubes (Fig. 3J, K). Moreover, pharmacological inhibition of m^6^A methylation using STM2457 in ALS myotubes reduced triglyceride levels in ALS myotubes (Fig. S4A-B). Collectively, these results demonstrate that METTL3-mediated m^6^A modification positively regulates triglyceride accumulation and that its downregulation contributes to impaired lipid metabolism in ALS skeletal muscle.

### METTL3-mediated m^6^A modification alleviates skeletal muscle atrophy in ALS by promoting triglyceride accumulation

Since triglycerides are a key reservoir for skeletal muscle energy metabolism ^[36, 37]^, we hypothesized that METTL3-mediated triglyceride accumulation would enhance energy supply in ALS myotubes. Consistent with this notion, ALS myotubes exhibited reduced fatty acid oxidation (FAO) levels and lower ATP content, both of which were significantly restored by *Mettl3* overexpression (Fig. 4A, B). Moreover, ALS myotubes displayed diminished survival under energy deprivation conditions, whereas *Mettl3* overexpression markedly improved cell viability (Fig. 4C). Notably, overexpression of *Mettl3^Mut^*had no significant effects (Fig. 4A-C). These results indicate that *Mettl3* overexpression enhances the cellular energy supply and resistance to energy stress.

**Figure 4.**
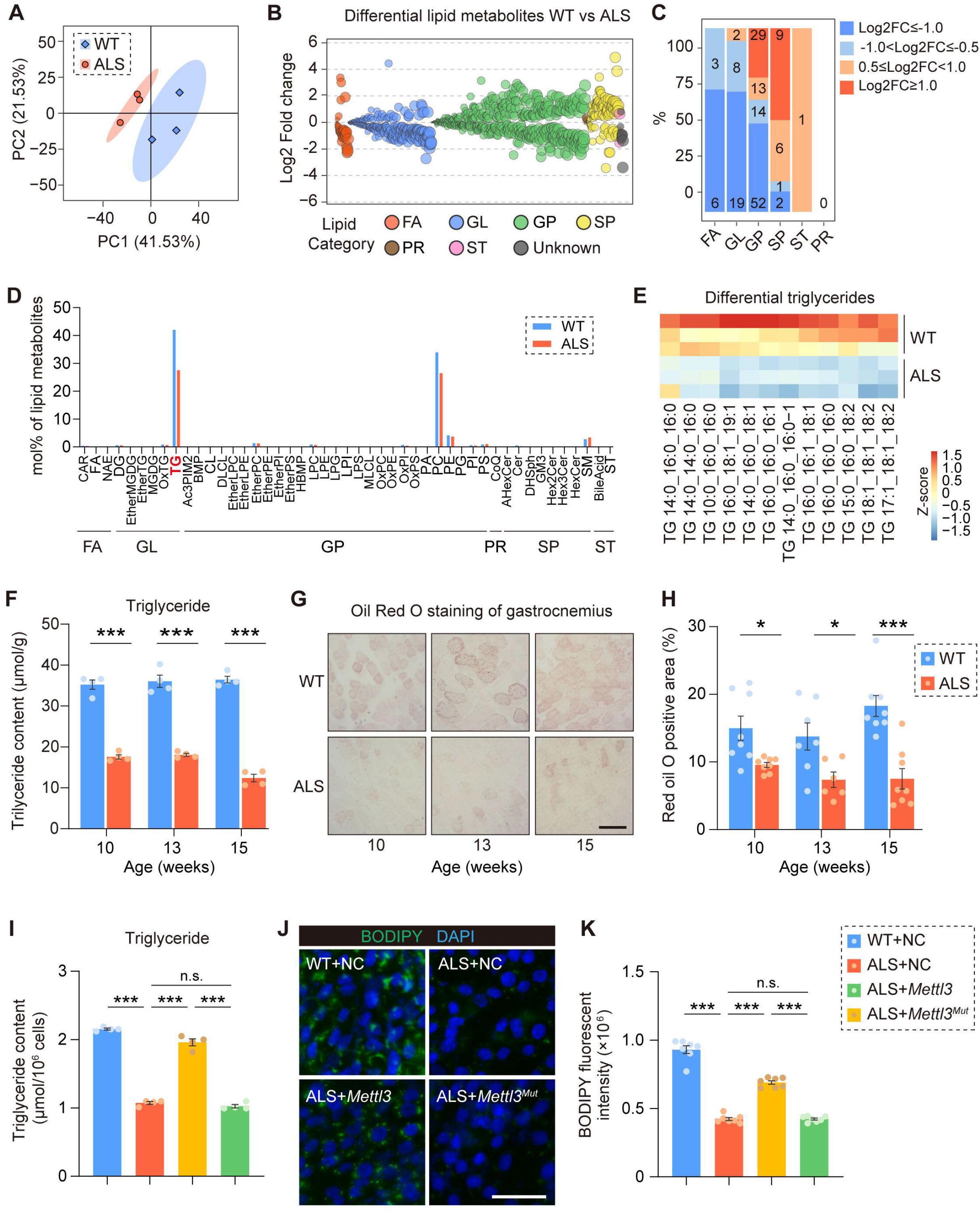
METTL3-mediated m^6^A modification alleviates skeletal muscle atrophy by promoting triglyceride accumulation. **(A)** Fatty acid oxidation activity detected by FAO blue staining in hSOD1^G93A^-tg ALS myotubes following *Mettl3* or its catalytic mutant overexpression (n = 8, 7, 8 and 7). **(B)** ATP levels in hSOD1^G93A^-tg ALS myotubes following *Mettl3* or its catalytic mutant overexpression (n = 3 per group). **(C)** Cell viability assessed by CCK-8 assay after 4 h of energy deprivation in hSOD1^G93A^-tg ALS myotubes following *Mettl3* or its catalytic mutant overexpression (n = 5, 5, 5 and 4). **(D)** Immunofluorescence staining of anti-MyHC in hSOD1^G93A^-tg ALS myotubes following overexpression of *Mettl3* or its catalytic mutant. MyHC, myosin heavy chain. Scale bar represents 100 μm. **(E)** Quantification of myotube diameter corresponding to panel **(D)** (n = 7 per group). **(F-I)** RT-qPCR analysis of *Atrogin-1* **(F)** (n = 4 per group), *MuRF-1* **(G)** (n = 4 per group), *MyoD* **(H)** (n = 4, 4 and 3) and *MyoG* **(I)** (n = 4 per group) mRNA expression in hSOD1^G93A^-tg ALS myotubes following *Mettl3* or its catalytic mutant overexpression. **(J)** Amplex red assay of triglyceride levels in hSOD1^G93A^-tg ALS myotubes following *Mettl3* or its catalytic mutant overexpression (n = 4 per group). **(K)** Fatty acid oxidation activity detected by FAO blue staining in hSOD1^G93A^-tg ALS myotubes after *Mettl3* overexpression with or without T863 treatment for 24h (n = 6, 7 and 7). **(L)** ATP levels in hSOD1^G93A^-tg ALS myotubes after *Mettl3* overexpression with or without T863 treatment for 24h (n = 3 per group). **(M)** Cell viability assessed by CCK-8 assay after 4 h of energy deprivation in hSOD1^G93A^-tg ALS myotubes after *Mettl3* overexpression with or without T863 treatment for 24h (n = 4 per group). **(N)** Immunofluorescence staining of anti-MyHC in *Mettl3* overexpressing hSOD1^G93A^-tg ALS myotubes with or without T863 treatment for 24h. Scale bar represents 100 μm. **(O)** Quantification of myotube diameter corresponding to panel **(N)** (n = 6 per group). **(P-S)** RT-qPCR analysis of *Atrogin-1* **(P)** (n = 4 per group), MuRF-1 **(Q)** (n = 3, 4 and 4), *MyoD* **(R)** (n = 4 per group) and *MyoG* **(S)** (n = 4 per group) mRNA expression in hSOD1^G93A^-tg ALS myotubes after *Mettl3* overexpression with or without T863 treatment for 24h. Data are presented as mean ± SEM. \**P* < 0.05, \*\**P* < 0.01, \*\*\**P* < 0.001; n.s., not significant.

We further investigated the role of METTL3 and m^6^A modification in ALS muscle atrophy. Results showed that *Mettl3* overexpression significantly restored the reduced myotube diameter observed in ALS myotubes (Fig. 4D, E). Consistently, the expression of atrophy-related biomarkers *Atrogin-1* (also known as *Fbxo32*) and *MuRF-1* (also known as *Trim63*) was decreased, whereas the expression of myogenic biomarkers *MyoG* and *MyoD* was increased following *Mettl3*, but not *Mettl3^Mut^*, overexpression (Fig. 4F-I). These results indicate that *Mettl3* overexpression alleviates ALS myotube atrophy. Conversely, pharmacological inhibition of m^6^A methylation using STM2457 further increased *Atrogin-1* and *MuRF-1* expression (Fig. S4C, D), indicating that loss of m^6^A modification exacerbates ALS myotube atrophy.

To determine whether the protective effects of METTL3 on ALS myotube atrophy depend on its regulation of triglyceride accumulation, we treated *Mettl3*-overexpressing cells with T863, a pharmacological inhibitor of triglyceride synthesis. T863 treatment abrogated the protective effects of *Mettl3* overexpression, as evidenced by reduced fatty acid oxidation, decreased ATP production, diminished resistance to energy stress (Fig. 4J-L). Consistently, treatment with T863 also resulted in reduced myotube diameter (Fig. 4M, N), increased expression of the atrophy markers *Atrogin-1* and *MuRF-1* (Fig. 4O, P), and decreased expression of the myogenic markers *MyoG* and *MyoD* (Fig. 4Q, R). These findings indicate that triglyceride availability is required for the protective effect of METTL3 against ALS myotube atrophy.

Taken together, these findings demonstrate that METTL3-mediated m^6^A modification enhances energy supply and protects ALS myotubes from atrophy by promoting triglyceride accumulation.

### Overexpression of METTL3 in skeletal muscle alleviates muscle atrophy and delays disease progression in ALS

To investigate whether targeting METTL3 could alleviate skeletal muscle atrophy and delay ALS progression *in vivo*, we overexpressed *Mettl3* in skeletal muscle by intravenous injection of adeno-associated virus 9 (AAV9) carrying the muscle-specific *Mck* promoter to WT and ALS mice at 4 weeks of age (Fig. 5A). Six weeks after *Mettl3*-overexpressing AAV administration, elevated METTL3 expression and increased global m^6^A levels were detected in skeletal muscle (Fig. 5B, C). No apparent histopathological abnormalities were observed in major organs, including the liver, lung, brain, kidney and spleen, indicating acceptable biosafety (Fig. S5).

**Figure 5.**
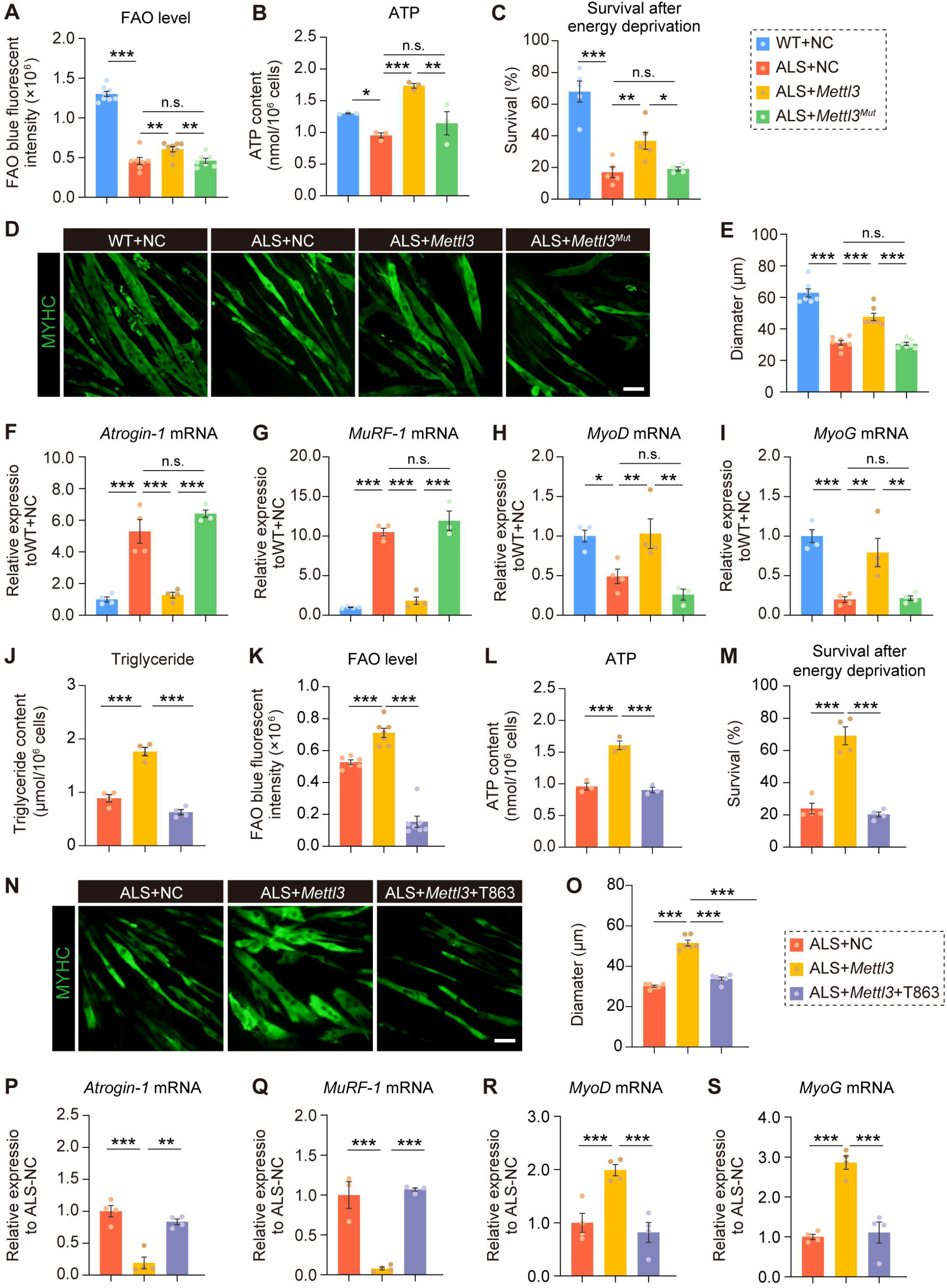
Overexpression of *Mettl3* in skeletal muscle promotes lipid accumulation, improves motor performance, and extends survival of ALS mice. **(A)** Schematic diagram of AAV-mediated *Mettl3* overexpression in the gastrocnemius of WT and hSOD1^G93A^-tg ALS mice. **(B)** Western blot analysis of METTL3 protein expression in gastrocnemius of WT and hSOD1^G93A^-tg ALS mice 6 weeks after *Mettl3*-overexpressing AAV administration (n = 3 per group). **(C)** Dot blot analysis of global m^6^A modification levels in gastrocnemius of WT and hSOD1^G93A^-tg ALS mice 6 weeks after *Mettl3*-overexpressing AAV administration (n = 3 per group). **(D)** Amplex red assay of triglyceride levels in gastrocnemius of WT and hSOD1^G93A^-tg ALS mice 6 weeks after *Mettl3*-overexpressing AAV administration (n = 4 per group). **(E)** Oil Red O staining of gastrocnemius of WT and hSOD1^G93A^-tg ALS mice 6 weeks after *Mettl3*-overexpressing AAV administration. Scale bar represents 100 μm. **(F)** Quantification of Oil Red O staining intensity from panel **(E)** (n = 6 per group). **(G-J)** RT-qPCR analysis of *Atrogin-1* **(G)** (n = 4 per group), *MuRF-1* (**H)** (n = 4 per group), MyoD **(I)** (n = 4 per group) and MyoG **(J)** (n = 4 per group) mRNA expression in gastrocnemius of WT and hSOD1^G93A^-tg ALS mice 6 weeks after *Mettl3*-overexpressing AAV administration. **(K)** Representative WGA staining of gastrocnemius from WT and hSOD1^G93A^-tg ALS mice 10 weeks after *Mettl3*-overexpressing AAV administration. Scale bar represents 100 μm. **(L)** Quantification of muscle fiber cross-sectional area from panel **(K)** (n = 6 per group). **(M)** Representative α-BTX labeling of neuromuscular junctions in the gastrocnemius of WT and hSOD1^G93A^-tg ALS mice 10 weeks after *Mettl3*-overexpressing AAV administration, followed by. Scale bar represents 100 μm. **(N)** Quantification of neuromuscular junction number from panel **(M)** (n = 4, 4, 3, and 4). Data are presented as mean ± SEM. \*\**P* < 0.01, \*\*\**P* < 0.001; n.s., not significant.

The triglyceride content in skeletal muscle was elevated following *Mettl3* overexpression (Fig. 5D). Oil Red O staining further confirmed enhanced neutral lipid accumulation (Fig. 5E, F), indicating that *Mettl3* overexpression promotes triglyceride accumulation *in vivo*.

Consistent with the *in vitro* experiments, the expression of the atrophy-related biomarkers *Atrogin-1* and *MuRF-1* was significantly downregulated and myogenic biomarkers *MyoD* and *MyoG* was significantly upregulated in the skeletal muscle of ALS mice following *Mettl3* overexpression (Fig. 5G-J). Morphological analysis revealed a significant increase in myofiber cross-sectional area in the skeletal muscle of ALS mice after *Mettl3* overexpression (Fig. 5K, L). Neuromuscular junction (NMJ) degeneration is a hallmark of skeletal muscle atrophy and dysfunction in ALS ^[3]^. Notably, α-bungarotoxin (α-BTX) labeling demonstrated that *Mettl3* overexpression partially restored NMJ numbers in ALS skeletal muscle (Fig. 5M, N). Collectively, these findings indicate that overexpression of *Mettl3* effectively alleviates skeletal muscle atrophy in ALS mice.

ALS mice typically exhibit body weight loss, motor dysfunction, and short lifespan ^[34, 38]^. Our results showed that overexpression of *Mettl3* in skeletal muscle alleviated body weight loss in ALS mice (Fig. 6A, B). Grip strength tests showed that *Mettl3* overexpression improved limb strength (Fig. 6C), and rotarod tests revealed significantly enhanced motor coordination (Fig. 6D). Gait analysis further indicated that *Mettl3* overexpression partially restored forelimb and hindlimb motor function at the age of 15 weeks (Fig. 6E, F). In addition, *Mettl3* overexpression significantly prolonged the survival of ALS mice (Fig. 6G). Therefore, these findings demonstrate that skeletal muscle-specific overexpression of *Mettl3* markedly alleviates motor impairment and extends lifespan in ALS mice.

**Figure 6.**
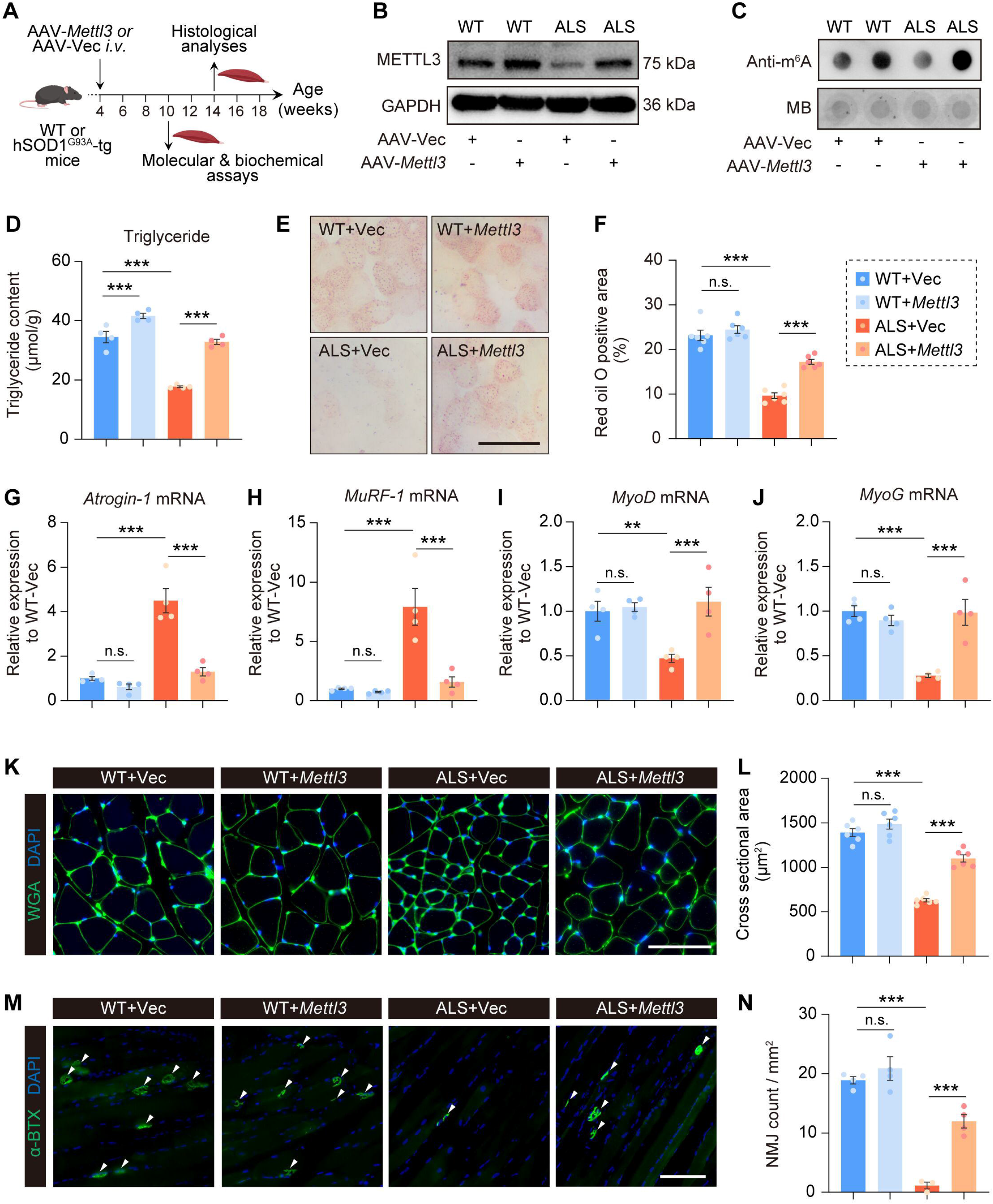
Overexpression of *Mettl3* in skeletal muscle alleviates ALS progression. **(A)** Schematic diagram of AAV-mediated *Mettl3* overexpression in the gastrocnemius of WT and hSOD1^G93A^-tg ALS mice. **(B)** Body weight analysis of WT and hSOD1^G93A^-tg ALS mice after *Mettl3* overexpression (n = 8, 10, 5 and 9). **(C)** Grip strength analysis of WT and hSOD1^G93A^-tg ALS mice after *Mettl3* overexpression (n = 5 per group). **(D)** Rota-rod analysis of WT and hSOD1^G93A^-tg ALS mice after *Mettl3* overexpression (n = 8, 10, 6 and 9). **(E)** Gait analysis based on footprint patterns of WT and hSOD1^G93A^-tg ALS mice after *Mettl3* overexpression (n = 3 per group). **(F)** Quantification of stride length corresponding to panel **(E)**. **(G)** Survival analysis of WT and hSOD1^G93A^-tg ALS mice after *Mettl3* overexpression (n = 8, 10, 6 and 9). Data are presented as mean ± SEM. \**P* < 0.05, \*\**P* < 0.01, \*\*\**P* < 0.001; n.s., not significant. For panels (**B-D)**, significance markers are indicated as follows: The upper symbols represent comparisons between WT+Vec and ALS+Vec groups, \**P* < 0.05, \*\**P* < 0.01, \*\*\**P* < 0.001. The lower symbols represent comparisons between ALS+Vec and ALS+*Mettl3* groups, #*P* < 0.05, ##*P* < 0.01, ###*P* < 0.001, n.s., not significant, u.d., undetectable.

### METTL3-mediated m^6^A modification enhances the mRNA stability of triglyceride metabolism-related genes

To identify downstream target genes of m^6^A modification involved in triglyceride accumulation, we performed an integrated analysis that combined three gene sets: (1) DEGs in skeletal muscle of ALS mice, (2) m^6^A-modified genes in mouse skeletal muscle, and (3) Lipid biosynthetic processes-related genes defined by GO annotations (Fig. 7A). This analysis yielded nine overlapping candidates. We then validated their expression and identified three genes, *Acsl1*, *Scd2*, and *Acaca*, that were significantly downregulated in the skeletal muscle of ALS mice (Fig. 7B). Among these, *Acsl1* encodes a critical enzyme responsible for fatty acid activation, a prerequisite step for glyceride synthesis, whereas *Acaca* and *Scd2* encode key enzymes essential for *de novo* fatty acid biosynthesis, thereby supporting glyceride production ^[39–41]^. (Fig. 7C)

**Figure 7.**
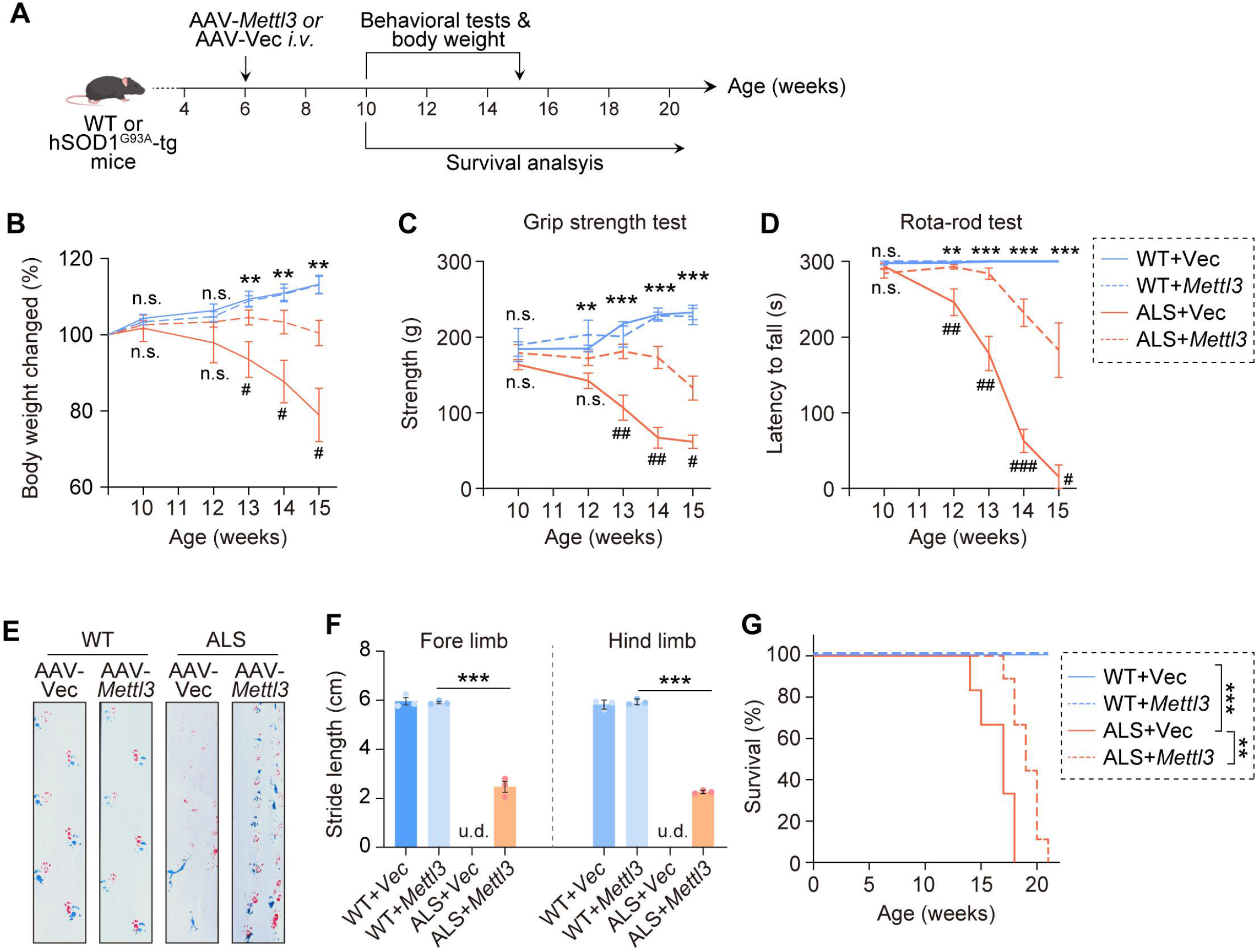
METTL3-mediated m^6^A modification enhances the expression of lipid metabolism-related genes by stabilizing their mRNA. **(A)** Venn diagram showing the overlap among DEGs in ALS mouse skeletal muscle (GEO accession: GSE245520), m^6T^A-modified genes in mouse skeletal muscle (GEO accession: GSE179368), and lipid biosythetic-related genes defined by the GO term (GO:0008610). **(B)** RT-qPCR analysis of mRNA expression of the overlapping genes identified in panel (**A**) in gastrocnemius of WT and hSOD1^G93A^-tg ALS mice at 15 weeks of age (n = 4 per group). **(C)** Schematic diagram illustrating the functions of *Acsl1*, *Scd2*, and *Acaca* in triglyceride metabolism. **(D-F)** IGV visualization of m^6^A modification peaks on the mRNAs of *Acsl1* **(D)**, *Scd2* **(E)**, and *Acaca* **(F)**. **(G-I)** m^6^A-MeRIP-qPCR analysis of m^6^A enrichment on *Acsl1* **(G)** (n = 4 per group), *Scd2* **(H)** (n = 3, 4, 4 and 4), and *Acaca* **(I)** (n = 4 per group) mRNAs in gastrocnemius of WT and hSOD1^G93A^-tg ALS mice at 15 weeks of age. **(J-L)** RT-qPCR analysis of *Acsl1* **(J)** (n = 4 per group), *Scd2* **(K)** (n = 4, 4, 4 and 3), and *Acaca* **(L)** (n = 4 per group) mRNA expression in hSOD1^G93A^-tg ALS myotubes following overexpression of *Mettl3* or its catalytic mutant. **(M-O)** RNA stability analysis of *Acsl1* **(M)** (n = 3 per group), *Scd2* **(N)** (n = 3 per group), and *Acaca* **(O)** (n = 3 per group) mRNAs in hSOD1^G93A^-tg ALS myotubes following *Mettl3* or its catalytic mutant overexpression after ActD (20 μg/ml) treatment. mRNA levels were quantified by RT-qPCR, and half-life values (t_1/2_) were calculated by linear regression. Data are presented as mean ± SEM. \*\**P* < 0.01, \*\*\**P* < 0.001; n.s., not significant, n.d. non-detectable.

Integrative Genomics Viewer (IGV) visualization revealed prominent m^6^A modification peaks near the stop-codon regions of *Acaca*, *Scd2*, and *Acsl1* (Fig. 7D-F). Moreover, m^6^A-MeRIP-qPCR demonstrated that m^6^A enrichment on these transcripts was markedly reduced in the skeletal muscle of ALS mice compared with WT mice (Fig. 7G-I).

To determine whether *Acsl1*, *Scd2*, and *Acaca* are regulated by METTL3-mediated m^6^A modification, we examined their expression following *Mettl3* overexpression *in vitro* and *in vivo*. All three genes were significantly downregulated in ALS myotubes compared with WT controls, but were restored upon *Mettl3*, instead of *Mettl3^Mut^*, overexpression (Fig. 7J-L). Consistently, *in vivo* overexpression of *Mettl3* in skeletal muscle also increased the expression of these genes (Fig. S6A-C).

Previous studies have shown that m^6^A modification regulates mRNA expression by modulating transcript stability ^[13, 14]^. Given that the mRNA levels of *Acsl1*, *Scd2*, and *Acaca* were upregulated upon *Mettl3* overexpression, we hypothesized that METTL3-mediated m^6^A modification enhances their stability. Consistent with this hypothesis, ActD (Actinomycin D) chase assay revealed that *Mettl3* overexpression attenuated the decay of *Acsl1*, *Scd2*, and *Acaca* mRNAs, whereas the catalytically inactive *Mettl3* mutant had no effect (Fig. 7 M-O).

Collectively, these results demonstrate that METTL3 promotes the expression of glyceride biosynthesis-related genes *Acsl1*, *Scd2*, and *Acaca* by stabilizing their mRNAs in an m^6^A-dependent manner.

### *Acsl1* is the primary downstream effector of METTL3-mediated triglyceride metabolism in ALS muscle

To identify the predominant downstream effector of METTL3-mediated triglyceride metabolism, we individually knocked down *Acsl1*, *Scd2*, and *Acaca* in *Mettl3*-overexpressing ALS myotubes. Knockdown of *Acsl1* almost completely abrogated the *Mettl3* overexpression-induced increase in triglyceride levels, whereas knockdown of *Scd2* or *Acaca* resulted in only modest reductions, revealing *Acsl1* as the primary downstream effector required for METTL3-mediated triglyceride accumulation (Fig. 8A). BODIPY staining further confirmed that knockdown of *Acsl1* significantly reduced triglyceride level mediated by *Mettl3* overexpression (Fig. 8B, C).

**Figure 8.**
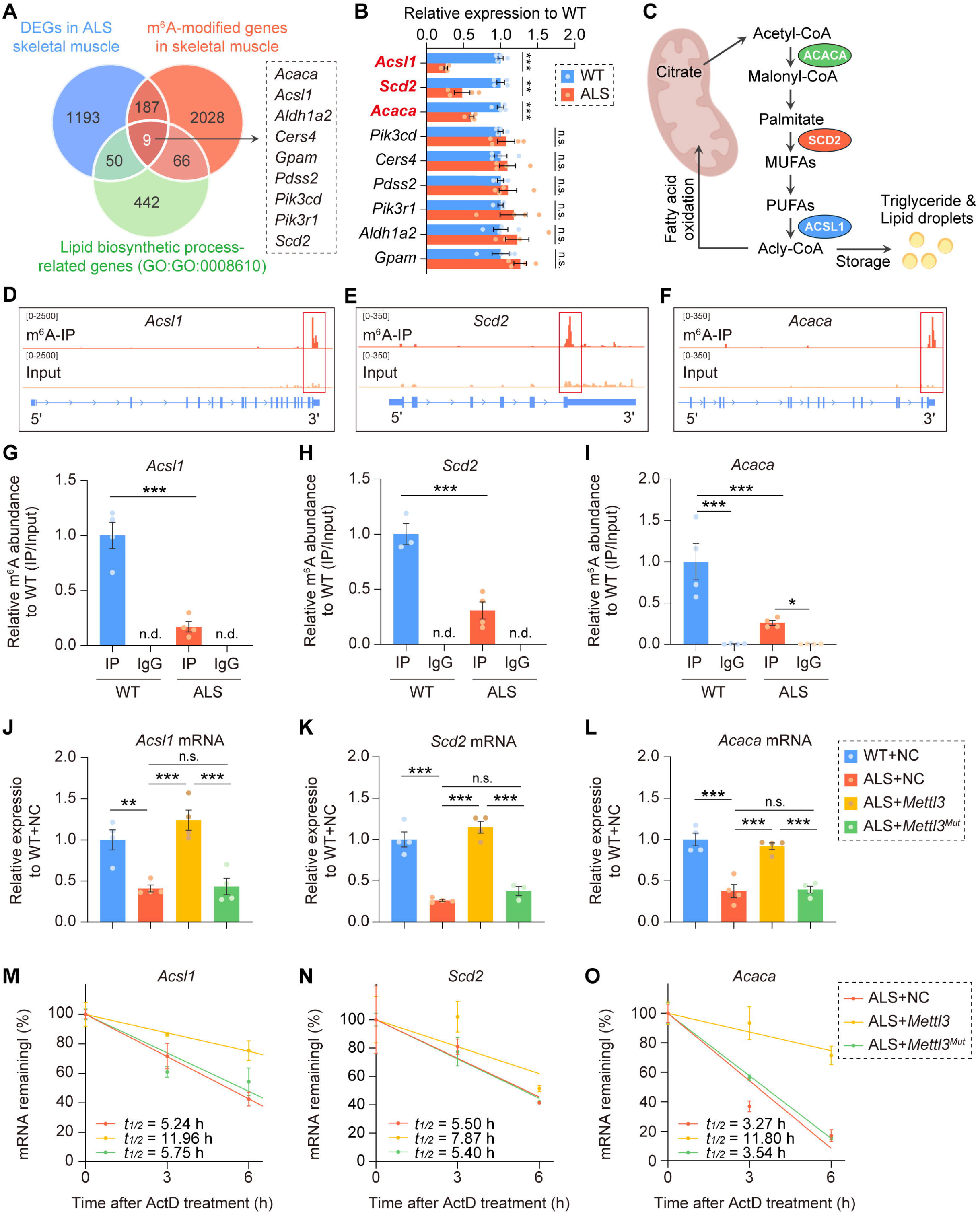
*Acsl1* knockdown abrogates the effects of *Mettl3* overexpression on triglyceride metabolism and muscle atrophy. **(A)** Amplex red assay of triglyceride levels in *Mettl3* overexpressing hSOD1^G93A^-tg ALS myotubes following *Acsl1*, *Acaca* or *Scd2* knockdown (n = 3 per group). **(B)** Representative BODIPY staining of *Mettl3* overexpressing hSOD1^G93A^-tg ALS myotubes following *Acsl1* knockdown. *siAcsl1*-1 and *siAcsl1*-2 represent two different siRNA sites. Scale bar represents 100 μm. **(C)** Quantification of BODIPY fluorescence intensity (n = 7 per group). **(D)** ATP levels in *Mettl3* overexpressing hSOD1^G93A^-tg ALS myotubes following *Acsl1* knockdown (n = 4 per group). **(E)** Cell viability assessed by CCK-8 assay after 4 h of energy deprivation in *Mettl3* overexpressing hSOD1^G93A^-tg ALS myotubes following *Acsl1* knockdown (n = 4 per group). **(F)** Immunofluorescence staining of anti-MyHC in *Mettl3* overexpressing hSOD1^G93A^-tg ALS myotubes following *Acsl1* knockdown. Scale bar represents 100 μm. **(G)** Quantification of myotube diameter corresponding to panel **(F)** (n = 6 per group). **(H-K)** RT-qPCR analysis of *Atrogin-1* **(H)** (n = 4 per group), *MuRF-1* **(I)** (n = 4 per group), *MyoD* **(J)** (n = 4 per group) and *MyoG* **(K)** (n = 4 per group) mRNA expression in *Mettl3* overexpressing hSOD1^G93A^-tg ALS myotubes following *Acsl1* knockdown. Data are presented as mean ± SEM. \*\**P* < 0.01, \*\*\**P* < 0.001; n.s., not significant, n.d. non-detectable.

Knockdown of *Acsl1* significantly reduced ATP content and impaired survival under energy deprivation in *Mettl3*-overexpressing ALS myotubes (Fig. 8D, E). Also, *Acsl1* knockdown decreased myotube diameter (Fig. 8F, G), increased the expression of the atrophy markers *Atrogin-1* and *MuRF-1*, and reduced the expression of the myogenic markers *MyoD* and *MyoG* (Fig. 8H-J).

Together, these findings establish *Acsl1* as the primary downstream effector required for METTL3-driven triglyceride accumulation and protective effects against ALS-associated skeletal muscle atrophy.

### IGF2BP2 is required for METTL3-mediated triglyceride metabolism and protective effects in ALS skeletal muscle

To identify the reader responsible for regulating *Acsl1* stability in ALS skeletal muscle, we individually knocked down candidate m^6^A readers known to modulate mRNA stability (*Igf2bp1/2/3* and *Ythdf2*) and found that only depletion of *Igf2bp2* significantly reduced *Acsl1* expression (Fig. 9A) ^[15]^. RIP-qPCR further demonstrated that IGF2BP2 directly binds to *Acsl1* mRNA (Fig. 9B). ActD chase assays showed that *Igf2bp2* knockdown abolished the stabilizing effect of *Mettl3* overexpression on *Acsl1* mRNA (Fig. 9C). These data indicate that METTL3-mediated stabilization of *Acsl1* is dependent on IGF2BP2.

**Figure 9.**
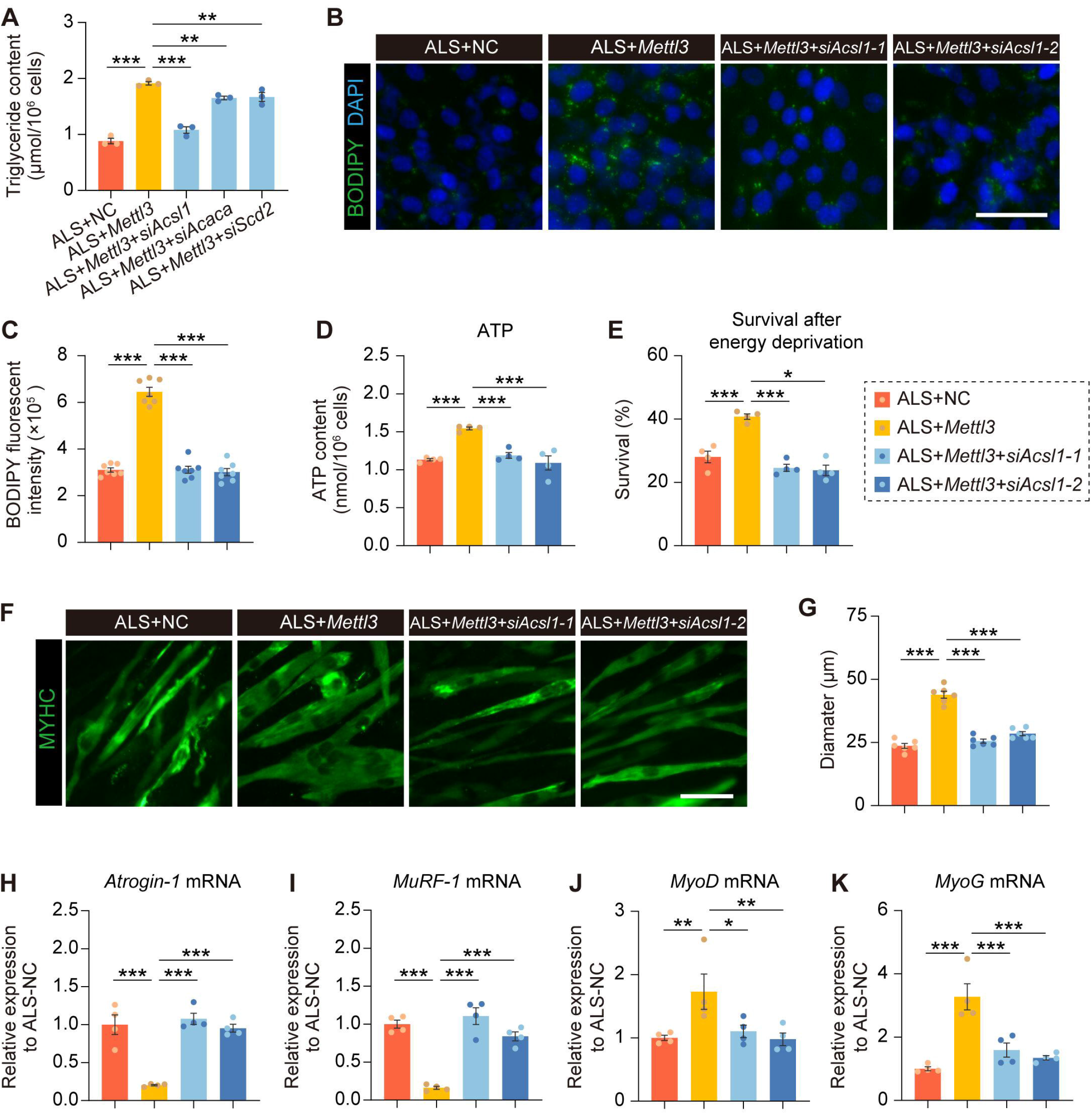
The regulatory effects of METTL3 on *Acsl1* expression, triglyceride metabolism and muscle atrophy are dependent on IGF2BP2. **(A)** RT-qPCR analysis of *Acsl1* mRNA expression in *Mettl3* overexpressing hSOD1^G93A^-tg ALS myotubes following different reader genes (*Igf2bp1/2/3* or *Ythdf2*) knockdown (n = 3 per group). **(B)** m^6^A-MeRIP-qPCR analysis of IGF2BP2 binding to *Acsl1* mRNA in the gastrocnemius of WT and hSOD1^G93A^-tg ALS mice at 15 weeks of age (n = 4 per group). **(C)** RNA stability analysis of *Acsl1* mRNAs in *Mettl3* overexpressing hSOD1^G93A^-tg ALS myotubes following *Igf2bp2* knockdown after ActD (20 μg/ml) treatment. mRNA levels were quantified by RT-qPCR, and half-life values (t_1/2_) were calculated by linear regression. **(D)** Amplex red assay of triglyceride levels in *Mettl3* overexpressing hSOD1^G93A^-tg ALS myotubes following *Igf2bp2* knockdown (n = 4 per group). *siIgf2bp2-1* and *siIgf2bp2-2* represent two different siRNA sites **(E)** Representative BODIPY staining of *Mettl3* overexpressing hSOD1^G93A^-tg ALS myotubes following *Igf2bp2* knockdown. Scale bar represents 100 μm. **(F)** Quantification of BODIPY fluorescence intensity (n = 7 per group). **(G)** ATP levels in *Mettl3* overexpressing hSOD1^G93A^-tg ALS myotubes following *Igf2bp2* knockdown (n = 4 per group). **(H)** Cell viability assessed by CCK-8 assay after 4 h of energy deprivation in *Mettl3* overexpressing hSOD1^G93A^-tg ALS myotubes following *Igf2bp2* knockdown (n = 4 per group). **(I)** Immunofluorescence staining of anti-MyHC in *Mettl3* overexpressing hSOD1^G93A^-tg ALS myotubes following *Igf2bp2* knockdown. Scale bar represents 100 μm. **(J)** Quantification of myotube diameter corresponding to panel **(I)** (n = 6 per group). **(K-N)** RT-qPCR analysis of *Atrogin-1* **(K)** (n = 4 per group), *MuRF-1* **(L)** (n = 3,4,4 and 4), *MyoD* **(M)** (n = 4 per group) and *MyoG* **(N)** (n = 4 per group) mRNA expression in *Mettl3* overexpressing hSOD1^G93A^-tg ALS myotubes following *Igf2bp2* knockdown. Data are presented as mean ± SEM. \*\**P* < 0.01, \*\*\**P* < 0.001; n.s., not significant, n.d. non-detectable.

To further determine whether IGF2BP2 is required for METTL3-mediated regulation of triglyceride metabolism and protective effects on muscle atrophy, we assessed the effects of *Igf2bp2* knockdown in *Mettl3*-overexpressing ALS myotubes. *Igf2bp2* depletion reversed the increase in triglyceride levels and neutral lipid accumulation induced by *Mettl3* overexpression (Fig. 9D-F). Consistently, *Igf2bp2* knockdown reduced ATP levels and impaired resistance to energy deprivation (Fig. 9G, H). Moreover, knockdown of *Igf2bp2* decreased myotube diameter (Fig. 9I, J), increased the expression of atrophy markers *Atrogin-1* and *MuRF-1*, and reduced the expression of myogenic markers *MyoD* and *MyoG* (Fig. 9K-N).

Collectively, our data demonstrate that METTL3 stabilizes *Acsl1* mRNA and protects ALS skeletal muscle in a IGF2BP2-dependent manner.

## Discussion

ALS has traditionally been viewed as a motor neuron-centric disorder, yet increasing evidence supports the “dying-back” hypothesis, which proposes that intrinsic defects within skeletal muscle actively contribute to disease initiation and progression ^[3, 4]^. In this study, we identify METTL3-mediated m^6^A modification as a key regulator of skeletal muscle homeostasis in ALS. Mechanistically, our data support a model in which METTL3-mediated m^6^A modification stabilizes *Acsl1* mRNA in a IGF2BP2-dependent manner, thereby sustaining triglyceride accumulation, preserving energy supply, and ultimately mitigating muscle atrophy. These findings establish an epitranscriptomic mechanism linking RNA modification to triglyceride metabolic homeostasis in ALS skeletal muscle.

Our findings provide mechanistic support for the “dying-back” hypothesis by demonstrating that intrinsic m^6^A modification dysregulation within skeletal muscle is sufficient to drive metabolic disorder and ALS progression. While previous studies have implicated muscle dysfunction in ALS, the upstream regulatory mechanisms have remained poorly defined. Here, our findings identify m^6^A RNA modification as an upstream regulator of triglyceride and energy metabolic remodeling, thereby identifying a previously unrecognized layer of muscle-intrinsic regulation in ALS pathogenesis.

Our lipidomics analysis identifies triglycerides as a central node of metabolic disruption in ALS skeletal muscle. Given their role as the principal intracellular energy reservoir, depletion of triglycerides are expected to reduce fatty acid oxidation and ATP production, thereby predisposing skeletal muscle to atrophy. Consistent with this notion, previous studies have shown that high-calorie or high-fat diets exert beneficial effects in ALS mouse models ^[42]^. Therefore, impaired triglyceride storage likely constitutes a key driver of energy deficiency and muscle atrophy while the restoration of triglyceride levels may serve as a feasible strategy to mitigate skeletal muscle atrophy in ALS.

Notably, however, a recent report described increased lipid content in the skeletal muscle of ALS patients, particularly in those carrying *FUS* mutations ^[43]^. These seemingly conflicting observations suggest that lipid metabolism in ALS is highly context-dependent and may be influenced by both genetic background and disease stage. Given that our study was limited to the hSOD1^G93A^ ALS mouse model, future studies integrating multi-omics approaches across different ALS subtypes (e.g., *C9ORF72*- or *FUS*- related ALS model) and disease stages will be essential to clarify the precise dynamics and effects of lipid metabolism in ALS pathogenesis.

Using integrated multi-omics analyses, we identified multiple genes involved in triglyceride biogenesis as downstream targets of METTL3, suggesting that METTL3 regulates triglyceride metabolism at multiple steps along the biosynthetic pathway. Among these candidates, *Acsl1*, which encodes a key enzyme responsible for fatty acid activation, emerged as the dominant functional effector mediating METTL3-dependent triglyceride regulation ^[39]^. In contrast, genes associated with *de novo* fatty acid synthesis, such as *Acaca* and *Scd2*, exhibited only modest contributions. This functional difference may be explained by the metabolic characteristics of skeletal muscle, where exogenous or stored fatty acids represent the predominant source for triglyceride synthesis, and *de novo* lipogenesis is relatively limited.

From a therapeutic perspective, our findings reveal muscle-specific overexpression of *Mettl3* using AAV vectors effectively ameliorated ALS-related phenotypes, suggesting the feasibility of targeting epitranscriptomic pathways *in vivo*. Given that neuron-specific *Mettl3* overexpression also exerts neuroprotective effects, as shown in our parallel unpublished studies, a combined strategy targeting m^6^A modification of both motor neurons and skeletal muscle may produce synergistic benefits. Future gene therapy approaches could exploit tissue-specific AAV vectors or small-molecule activators of METTL3 to achieve coordinated restoration of m^6^A homeostasis across the neuromuscular axis.

In summary, our study identifies METTL3-mediated m^6^A modification as a critical regulator of skeletal muscle metabolism in ALS and establishes a mechanistic link between RNA methylation, lipid metabolism, and muscle atrophy (Fig. 10). By uncovering an epitranscriptomic pathway that governs metabolic homeostasis, these findings provide a conceptual framework for understanding muscle-intrinsic contributions to ALS and open new avenues for therapeutic intervention.

**Figure 10.**
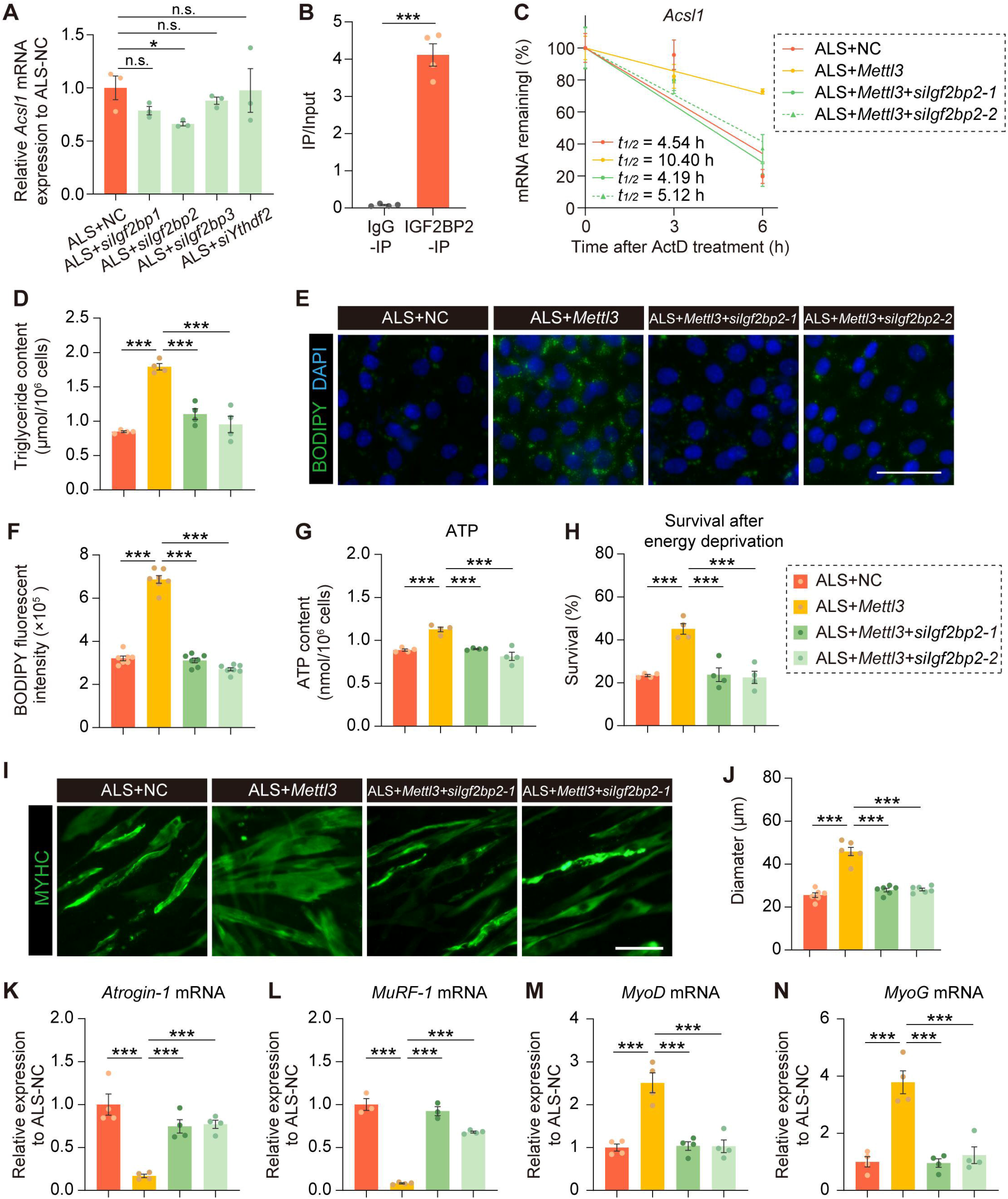
Proposed model In ALS skeletal muscle, reduced METTL3-mediated m^6^A modification leads to the downregulation of lipid metabolism-related genes, particularly *Acsl1*, resulting in impaired triglyceride accumulation and insufficient energy supply, ultimately contributing to muscle atrophy. Mechanistically, IGF2BP2 functions as the m^6^A reader that promotes *Acsl1* mRNA stability, thereby mediating the regulatory effects of m^6^A modification on *Acsl1* expression.

## Acknowledgements

This work was supported by the National Natural Science Foundation of China (82271483 and 82571627), Shandong Provincial Natural Science Foundation (ZR2024QH628 and ZR2024MH112), and Weifang Science and Technology Development Plan Project (2024YX033).

We would like to thank the Intelligent Medical Engineering Experimental Center, School of Basic Medical Sciences, Shandong Second Medical University, for providing the server platform.

## Data Availability

Lipidomics data generated in this study have been deposited in the OMIX, China National Center for Bioinformation / Beijing Institute of Genomics, Chinese Academy of Sciences, project ID: PRJCA050591.

