## Supplemental Figures S1-S6 for "METTL3-mediated m^6^A modification alleviates skeletal muscle atrophy in ALS by regulating ACSL1-dependent triglyceride metabolism"

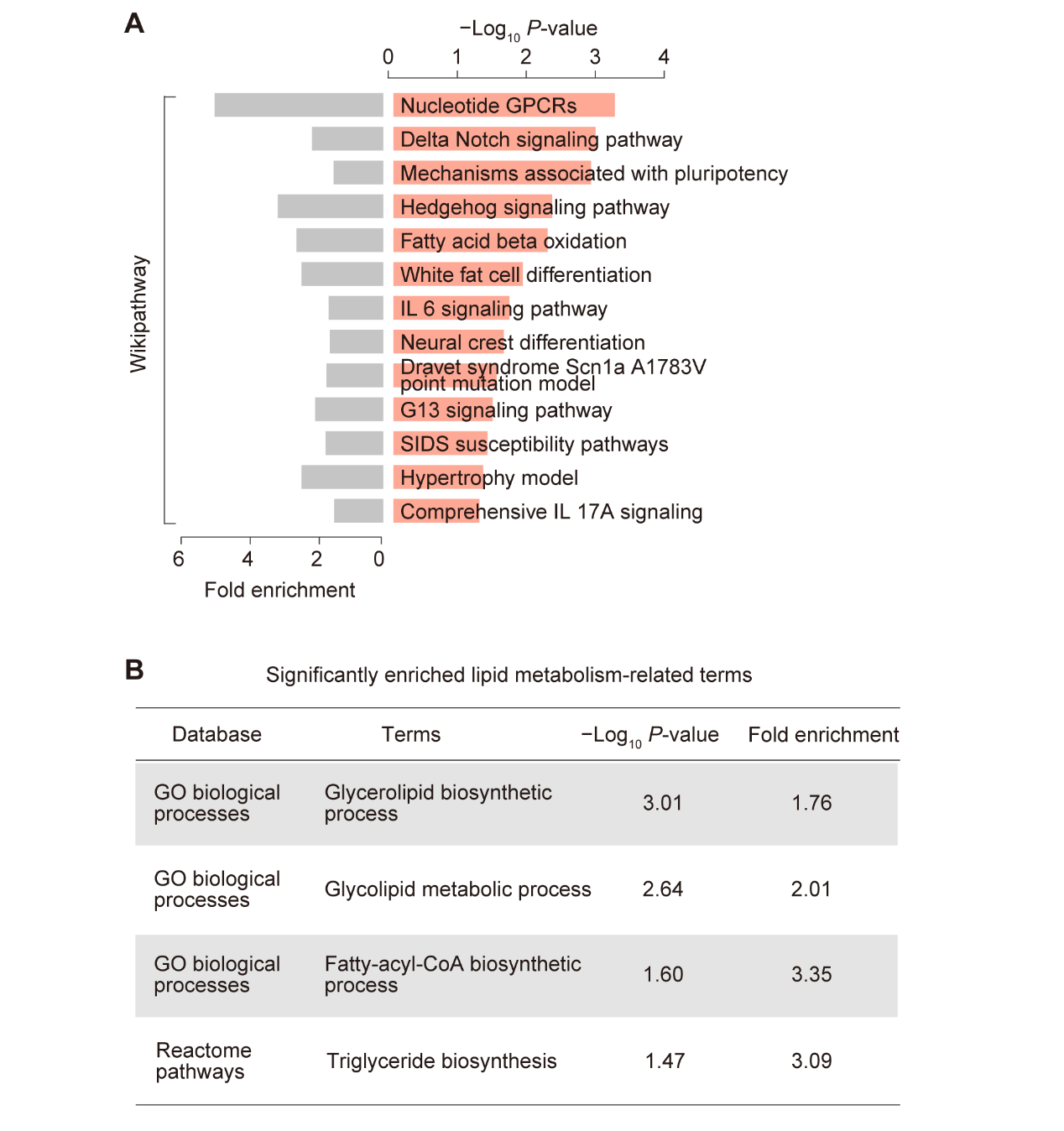

Figure S1. m^6^A-modified genes in mouse skeletal muscle are functionally enriched in lipid metabolism-related pathways..

**(A)** Wikipathway enrichment analysis of m^6^A-modified genes identified in mouse skeletal muscle.

**(B)** Significantly enriched lipid metabolism-related terms of m^6^A-modified genes in mouse skeletal muscle. The full enrichment results are provided in the Supplementary Data 2.

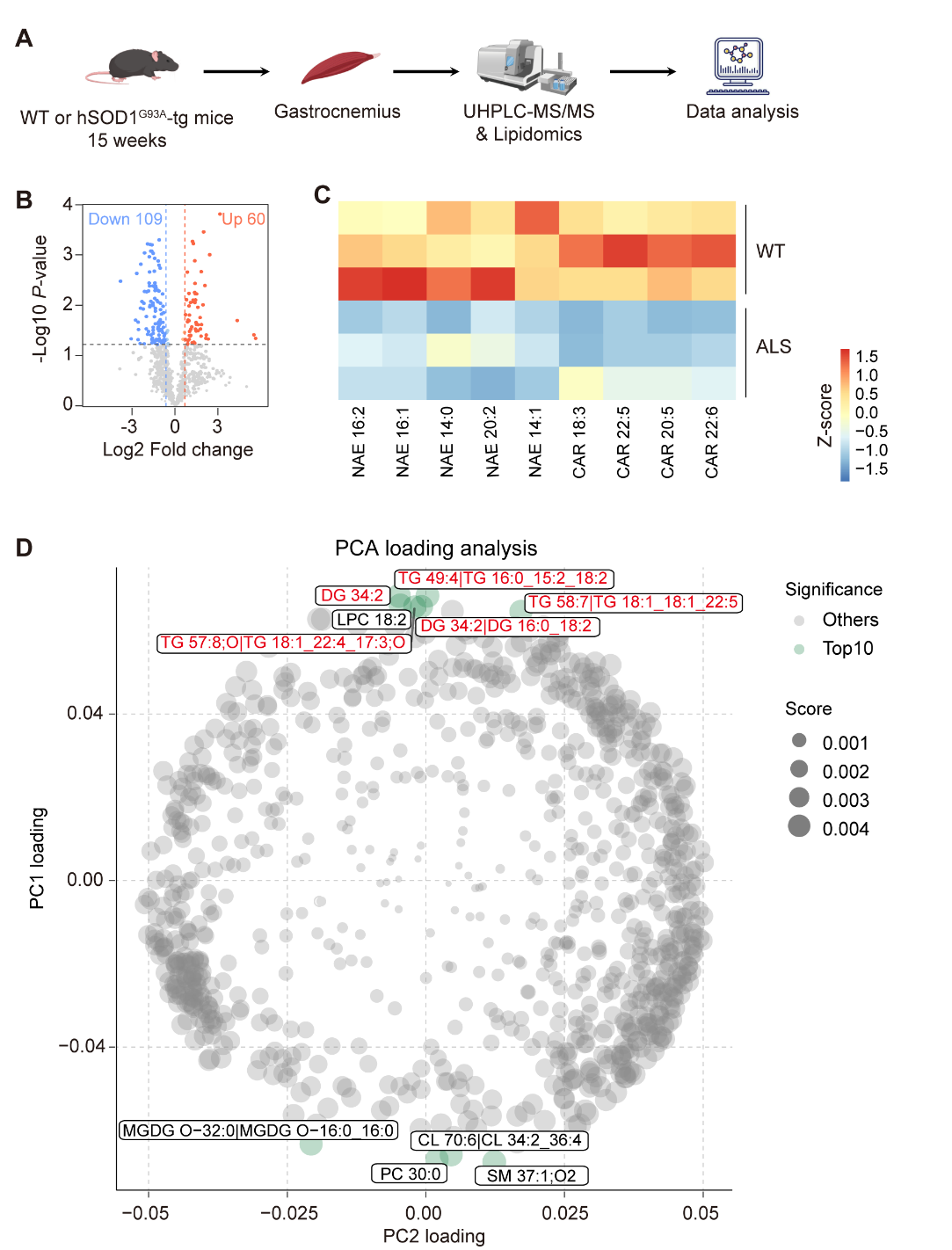

Figure S2. Lipidomics analysis reveals dysregulated fatty acid and glyceride levels in the skeletal muscle of hSOD1^G93A^-tg ALS mice.

**(A)** Schematic overview of the lipidomics analysis workflow.

**(B)** Volcano plot showing differential lipid metabolites between WT and hSOD1^G93A^-tg ALS gastrocnemius. Differential metabolites were defined by Log_2_ fold change > 1.5 and *P* < 0.05.

**(C)** Heatmap displaying changes in fatty acyls between the gastrocnemius of WT and hSOD1^G93A^-tg ALS mice
**(D)** PCA loading analysis of lipid metabolites contributing to principal components. Each point represents an individual lipid metabolite, with point size proportional to its contribution score. Metabolites ranked among the top 10 contributors are highlighted in green, while the symbol of triglyceride species are labeled in red.

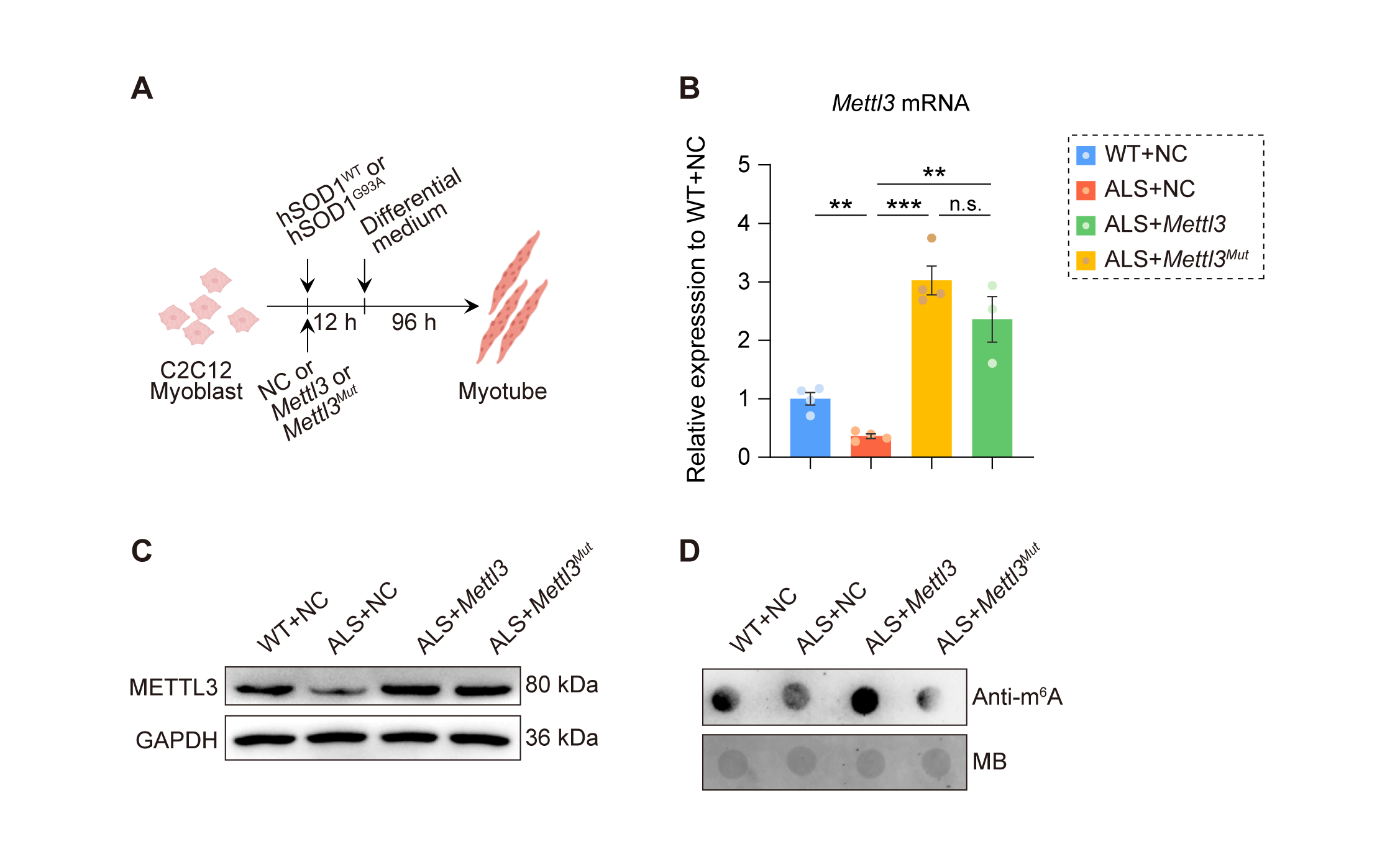

Figure S3. Validation of *Mettl3* overexpression in the hSOD1^G93A^-tg ALS myotube model.

**(A)** Schematic overview of the *in vitro* cultured ALS myotube model generated by differentiating hSOD1^G93A^-overexpressing C2C12 myoblasts, followed by *Mettl3* or catalytic mutant *Mettl3* overexpression.

**(B)** RT-qPCR analysis of *Mettl3* mRNA expression in hSOD1^G93A^-tg ALS myotubes following overexpression of *Mettl3* or its catalytic mutant (n = 4, 4, 4 and 3).
**(C)** Western blot analysis of METTL3 protein expression in hSOD1^G93A^-tg ALS myotubes following overexpression of *Mettl3* or its catalytic mutant (n = 3 per group).
**(D)** Dot blot analysis of m^6^A modification levels in hSOD1^G93A^-tg ALS myotubes following overexpression of *Mettl3* or its catalytic mutant (n = 3 per group).

Data are presented as mean ± SEM. ***P* < 0.01, ****P* < 0.001; n.s., not significant.

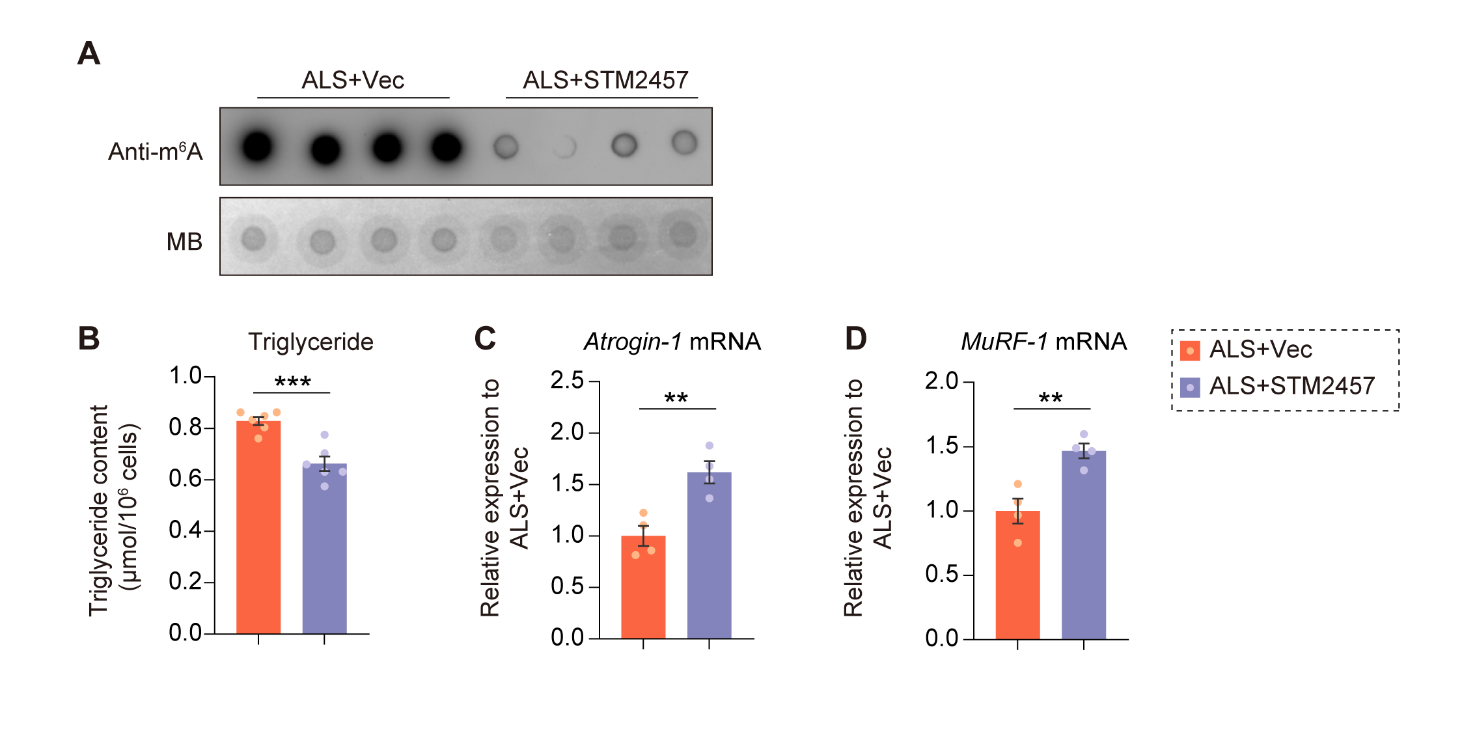

Figure S4. Pharmacological inhibition of m^6^A formation reduces lipid accumulation and increases atrophy biomarker expression.

**A)** Dot blot analysis of m^6^A modification levels showing the inhibitory efficiency of STM2457 (25 μM).

**(B)** Amplex red assay of triglyceride levels (n = 6 per group) in hSOD1^G93A^-tg ALS myotubes after 24 h of STM2457 treatment (25 μM).

**(C-D)** RT-qPCR analysis of *Atrogin-1* (**C**) (n = 4 per group) and *MuRF-1* (**D**) (n = 4 per group) mRNA expression in hSOD1^G93A^-tg ALS myotubes after 24 h of STM2457 treatment (25 μM).

Data are presented as mean ± SEM. ***P* < 0.01, ****P* < 0.001; n.s., not significant.

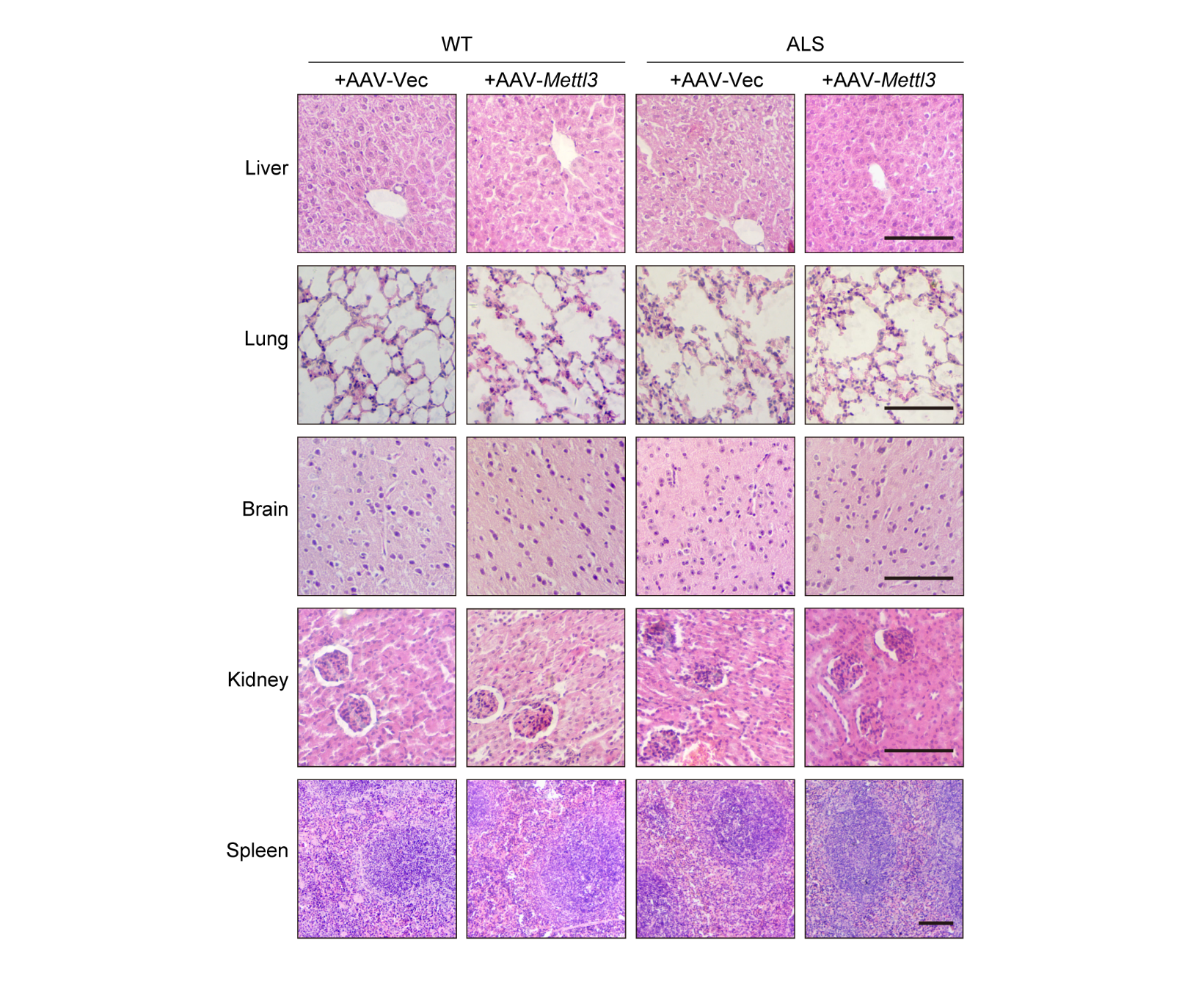

Figure. S5 H&E staining of major tissues from hSOD1^G93A^-tg ALS mice following *in vivo* *Mettl3* overexpression.

Representative H&E staining images of the liver, lung, brain, kidney, and spleen from WT and hSOD1^G93A^-tg ALS mice 6 weeks after AAV-mediated *Mettl3* overexpression. Scale bar represents 100 μm.

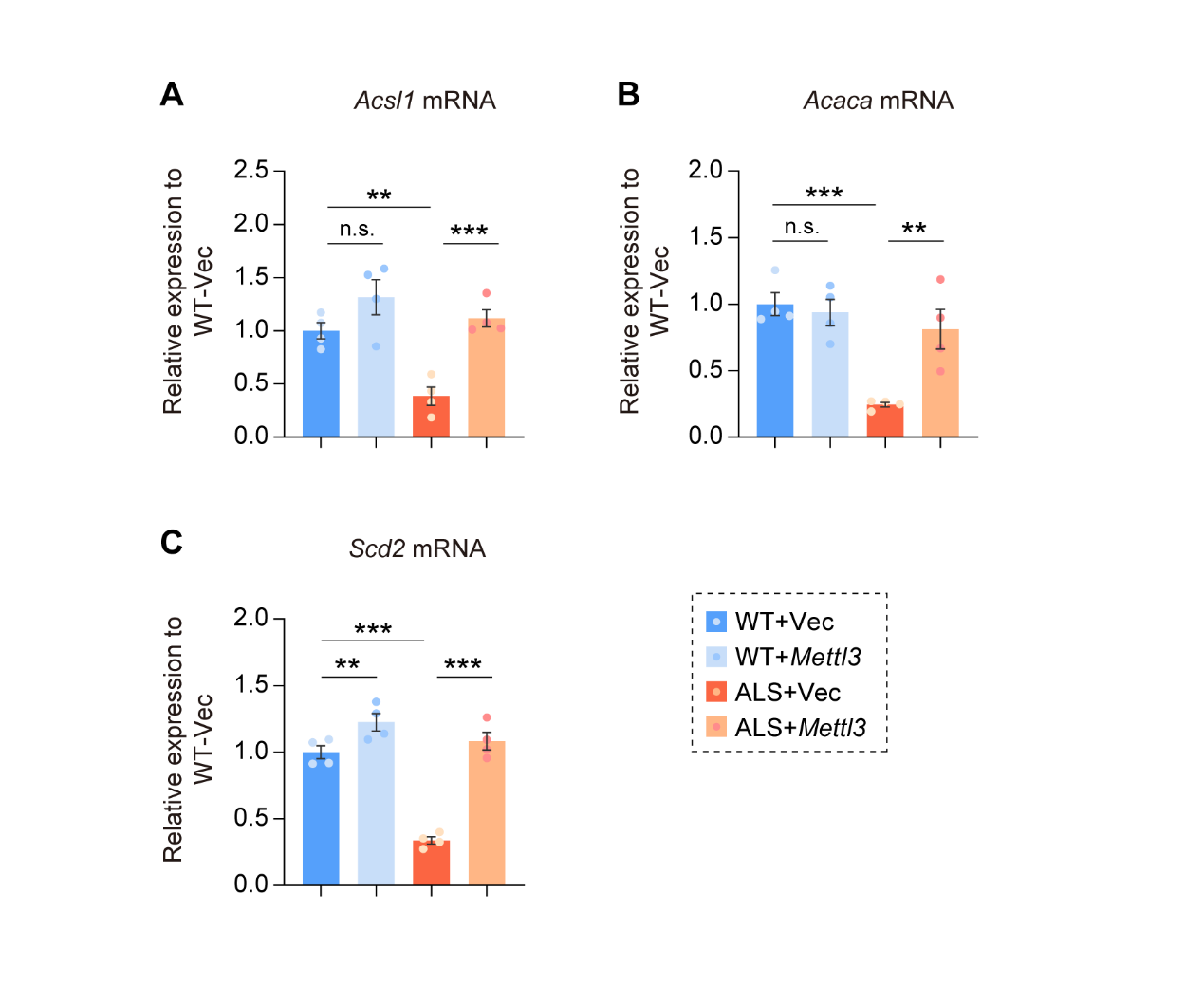

Figure. S6 Overexpression of *Mettl3* in skeletal muscle of hSOD1^G93A^-tg mice promote the epxression of *Acsl1*, *Acaca* and *Scd2*.

**(A-C)** RT-qPCR analysis of *Acsl1* (A) (n = 4 per group), *Acaca* (B) (n = 4 per group) and *Scd2* (C) (n = 4 per group). in the gastrocnemius of WT and hSOD1^G93A^-tg ALS mice 6 weeks after AAV-mediated *Mettl3* overexpression.

Data are presented as mean ± SEM. ***P* < 0.01, ****P* < 0.001; n.s., not significant.
